# A maternal effect regulates global DNA methylation patterns

**DOI:** 10.1101/2020.08.15.252239

**Authors:** Remco Loos, Valeria Carola, Enrica Audero, Elena Brini, Luisa Lo Iacono, Anna Moles, Paul Bertone, Cornelius Gross

## Abstract

Variation in DNA methylation between individuals has been shown to be influenced by both genetic and environmental factors. However, the relative impact of genetic and non-genetic factors on DNA methylation patterns across the mammalian genome has not been systematically studied. We performed whole-genome methylation analysis in two inbred mouse strains, revealing striking differences in the global distribution of DNA methylation. Although global methylation patterns were indistinguishable for most genomic features, a significant increase in the number of unmethylated CpG-island promoters and first exons was observed between strains. Experiments using F1 reciprocal hybrid strains demonstrated that the genotype of the mother dictated global DNA methylation patterns. Cross-fostering experiments ruled out a postnatal maternal effect on these differences and suggested that they were driven by a prenatal maternal effect, possibly via differential deposition of maternal gene products into the oocyte or uterine environment. These data demonstrate that maternal effects have a major impact on global DNA methylation patterns.

## Introduction

DNA cytosine methylation is a long-lasting epigenetic modification occurring primarily in the context of the CpG dinucleotide in somatic cells. Specifically, it has been established that methylated promoter regions can repress transcription of the associated gene (Jaenisch and Bird, 2003), suggesting a critical role for the DNA methylation profile of a cell in establishing its transcriptional activity. Methylation patterns are known to vary across cell types, where they contribute to the maintenance of cellular identities and to the response to environmental perturbations through the regulation of gene expression (Rakyan et al., 2008;Lister et al., 2009). In the brain, DNA methylation status has been associated with its structure and function (Wheater et al., 2020) and aberrations in methylation profiles, with sub-regional and cell-type specificity (Rizzardi et al., 2019) have been associated with many forms of mental illness (Grayson and Guidotti, 2013;Shimada-Sugimoto et al., 2017;Liu et al., 2018;Li et al., 2019).

The mechanisms by which DNA methylation patterns are established during development and maintained or remodelled during adulthood are not well understood, but differences between individuals can be linked to both genetic and non-genetic factors. Although studies in mouse and human found that the majority of observed differential methylation could be attributed to genetic differences (Schilling et al., 2009;Gertz et al., 2011), specific prenatal and postnatal environmental factors, including maternal nutrition, quality of maternal care, season of conception, and exposure to chemicals (Weaver et al., 2004;Waterland et al., 2010;Banik et al., 2017;Arsenault et al., 2018;Lester et al., 2018) have been shown to influence DNA methylation. Indeed, epigenetic processes, and DNA methylation in particular, have been proposed to mediate the interaction between genetic predisposition and environmental pressure, contributing to brain development and diseases. This sort of epigenetic plasticity may be restricted to specific developmental time windows, environmental factors, tissues and genomic regions. However, this phenomenon has not been systematically studied, and the relative contributions of genetic and non-genetic factors to individual variation in DNA methylation remain to be investigated.

Here we report an unbiased, whole-genome, high-resolution DNA methylation mapping of hippocampal DNA from two widely used inbred mouse strains and their reciprocal F1 hybrids. Genome-wide methylation profiles were generated using the Methylation Mapping Analysis by Paired-end Sequencing (Methyl-MAPS) method (Edwards et al., 2010) on the Applied Biosystems SOLiD sequencing platform. These profiles allow for a relatively high-resolution genome-wide study of individual differences in DNA methylation. Moreover, the controlled experimental set-up makes it possible to disentangle genetic and environmental influences on global methylation patterns.

## Methods

### Animals and tissue preparation

C57BL/6J@Ico (C57BL/6) and BALB/cByJ@Ico (BALB/c) mice were purchased from Charles River Laboratories (Calco, Italy). C57BL/6 and BALB/c strains were produced through inbreeding. Reciprocal F1 hybrid mice were obtained by breeding C57BL/6 females and BALB/c males (C57BL/6×BALB/c) and BALB/c females and C57BL/6 males (BALB/c×C57BL/6). For the production of F1 hybrid (BALB/c×C57BL/6) cross-fostered animals, BALB/c females were mated with C57BL/6 males and F1 offspring were cross-fostered within 1 day of parturition to either a BALB/c or C57BL/6 mother that had given birth to her own litter on the same day. All BALB/c and C57BL/6 foster mothers had been previously bred with C57BL/6 and BALB/c male mice, respectively.

Matings were set up at 10 weeks of age. For all breeding conditions, fathers were removed before parturition and mothers and offspring were left undisturbed until postnatal day 21, at which time pups were weaned and housed between three and five per cage and weekly cage cleaning was resumed. No effort was made to normalize litter sex composition or size. Food and water were provided ad libitum, and mice were housed on a 12:12 light:dark cycle with lights on at 07:00 AM. Adult males were sacrificed by decapitation at 8-10 weeks of age. Both hippocampal hemispheres were rapidly dissected, frozen on dry ice, and stored at −80 °C until the tissue was processed. Animals were handled in strict accordance with good animal practice as defined by the relevant national and local animal welfare bodies. The experiments for this study were approved by the ethics committee of the Italian Department of Health and conducted under license/approval ID #:91/2007-B, according to Italian regulations on the use of animals for research (legislation DL 116/92) and NIH guidelines on animal care.

### Isolation of genomic DNA and enzymatic fractionation

Genomic DNA was collected from the right hippocampus of adult male mice. DNA was prepared by tissue fragmentation and overnight digestion at 55°C in 10 mM NaCl, 10 mM Tris-HCl pH 8.0, 25 mM EDTA, 0.5% SDS, 100 ng/mL proteinase K. DNA was purified by three rounds of phenol/chloroform extraction followed by RNase A treatment and ethanol precipitation. DNA quality was confirmed by agarose gel electrophoresis and quantification was performed on a NanoDrop 8000 spectrophotometer (Thermo Scientific).

Enzymatic digestion was performed using five tetranucleotide methylation-sensitive restriction endonucleases (henceforth RE) on half of the DNA, and the methylation-dependent McrBC complex on the other half. The RE enzymes cut DNA molecules between the C and G nucleotides of their recognition sites (AciI: CCGC, BstUI: CGCG, HhaI: GCGC, HpaII: CCGG, HpyCH4IV: ACGT), while the McrBC complex recognizes two G/A^m^C sites between 40bp and 3kb apart, with no methylated C nucleotides between these sites. McrBC cleaves DNA at locations approximately 25-35 bp inside the recognition sites.

For each experimental group, DNA samples from 8 different specimens were pooled in equimolar concentration to a total of 12 μg. Limited digestion of 6 μg DNA with McrBC and 6 μg DNA with RE was performed, as described previously (Edwards et al., 2010).

### Library construction and high-throughput sequencing

Paired-end libraries were prepared from the DNA fractions digested with RE and McrBC as previously described (Edwards et al., 2010). Libraries were subjected to mate-pair sequencing on the Applied Biosystems SOLiD System DNA sequencer to obtain 2×30bp reads for the C57BL/6, BALB/c and cross-fostered BALB/c×C57BL/6 samples, and 2×25bp reads for C57BL/6×BALB/c and BALB/c×C57BL/6 samples. Sequencing yielded between 52 and 89 million paired reads per library, amounting to 107 to 175 million read pairs per biological sample (considering two digestion libraries for each sample). Further details can be found in **Supplemental Table 5**. Sequencing files have been deposited in the Sequence Read Archive under the study accession number ERP000909 (European Nucleotide Archive).

### Alignment and filtering

The 30 bp reads were trimmed to 27 bp, since any further bases represent adapter sequence. Sequences were aligned to the July 2007 assembly of the mouse genome (NCBI37/mm9) using MAQ (Li et al., 2008), with default settings regarding mismatches and a maximum insert size of 20 kb. Only the reads that were successfully aligned as a pair were retained and used to infer the coordinates of the original RE and McrBC fragments. Recognition sites for all enzymes in the genome were identified with a custom Perl script, using the Ensembl API (Flicek et al., 2011). Fragments were then filtered using the following criteria, similar to those used previously (Edwards et al., 2010):

1. **RE fragments**: a RE recognition site must be present within 10 bp of at least one end of the fragment
2. **McrBC fragments**: an McrBC recognition site must be present within 50 bp of at least one end of the fragment

The numbers of filtered fragments for each sample are given in **Supplemental Table 5**.

### Inference of methylation status

The methylation status of each CpG interrogated by both methods (that is, contained in a recognition site of one of the RE enzymes and of the McrBC enzyme) was determined by combining the evidence provided by the two enzymatic digestions. The interpretation of fragment coverage in terms of methylation status of a given CpG is as follows:

1. If a CpG is covered by a RE fragment, this means the CpG was not cut by the methylation sensitive enzymes and thus is *methylated*
2. If a CpG borders a RE fragment, this CpG was cut by the methylation sensitive enzymes and thus is *not methylated*
3. If a CpG is well inside an McrBC fragment (>50 bp), the CpG was not cut by the enzyme and thus is *not methylated*

For a given CpG we can find evidence of both methylation and non-methylation, reflecting allele-specific differences or heterogeneity in the tissue sample. The methylation status is represented by a score between 0 and 1, where 0 represents a non-methylated CpG and 1 a fully methylated CpG. However, since the evidence of the two enzymatic digestions is not directly comparable, normalization is required before computing the methylation score. The numbers of fragments obtained for each treatment is determined by technical issues such as sequencing depth and does not necessarily reflect the proportion found in the samples.

We used the proportion of genome coverage of the McrBC and RE samples to correct the fragment counts. If the fragments obtained from one digestion cover a larger fraction of the genome, one would expect to find a larger proportion of these fragments if their number were not bounded artificially by sequencing depth. Computing the genome coverage of random subsets of the data shows that our dataset is highly saturated with respect to genome coverage, with half of the fragments providing over 98% of the coverage of the whole set (**Supplemental Fig. 2**). This means that the coverage we find is a good approximation of the actual genome coverage we expect from the fragments obtained by each enzymatic digestion.

The correction factor, which also takes into account the number and size of fragments in each sample, is calculated as follows:

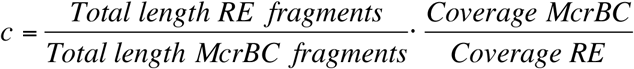

This factor is applied to the counts of the McrBC fragments. The methylation score *m* for a CpG is then defined as:

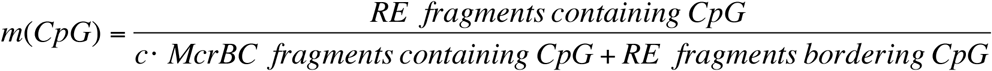

A minimal coverage of four fragments is required to assign a methylation score to a CpG.

### Methylation analysis for genomic elements

Data for annotated CpG islands, Ensembl genes and repeat regions were downloaded from Ensembl release 60 (Flicek et al., 2011). Gene promoters are defined here as the region 2 kb upstream and 200 bp downstream of the annotated transcription start site. Promoters are considered CpG island promoters if a part of the promoter region intersects with a CpG island. Similarly, promoter CpG islands are those CpG islands which intersect with a promoter region.

The average methylation score for a genomic element *e* is computed as:

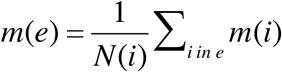

where *i* denotes an individual CpG and *N*(*i*) is the number of scored CpGs in the region. This score is computed only for elements which contain at least five scored CpGs, and for which more than half of the interrogated CpGs have sufficient coverage to call methylation status.

### Differential methylation and functional annotation

Differentially methylated regions (DMRs) were identified using a sliding window approach. Large DMRs were defined as regions at least 800 bp in length, contain more than 15 scored CpGs, and having a difference in average methylation of >0.3, with the largest CpG-level difference >0.5. A large DMR was associated with a gene if it was located either in the gene body or within a defined promoter-proximal region (<5kb upstream from TSS). Gene Ontology (GO) enrichment analysis was performed using the GOstats R package (Falcon and Gentleman, 2007) and DAVID (Huang da et al., 2009a;b). P-values were corrected for multiple testing using the Benjamini & Hochberg correction. The reported P-values are the more conservative set obtained from DAVID.

Unsupervised hierarchical clustering and heatmap plotting were performed in R. For the whole-genome clustering, the methylation scores for 1000 bp bins over the entire genome were computed in the same manner as described above for genomic elements. The clustering and heatmap were based on those bins that showed a difference in methylation score of >0.6 between the minimum and maximum value across the four samples. The clustering and heatmaps of CpG islands and CpG island promoters comprise regions displaying a difference in methylation score of >0.4 between the minimum and maximum value across the four samples.

## Results

We first investigated DNA methylation status in adult hippocampal tissues of two widely used inbred mouse strains, C57BL/6 and BALB/c. Methyl-MAPS profiling revealed the methylation status of 5.5 million CpGs in each sample, capturing over 98% of all genomic CpGs containing restriction sites compatible with the method and over 26% of all CpGs present throughout the genome.

In the methyl-MAPS method, two complementary enzymatic digestions are used to infer the methylation state of a given CpG, expressed as a score between 0 and 1, with 1 being fully methylated (see Methods). A considerable number of all profiled CpGs, 54.2% in C57BL/6 and 45.7% in BALB/c, were found to be in a partially methylated state (methylation score between 0.3 and 0.7). A total of 1,068 large (> 800 kb) differentially-methylated regions (DMRs) throughout the genome were identified between the two strains. Functional analysis of these DMRs and their proximal genes showed significant enrichment in categories related to embryonic development, chromatin organization, and transcriptional regulation (**Supplementary Tables 1,2**).

To determine the relationship between DNA methylation variation and functional elements in the genome, we examined the degree of methylation in exons, introns, CpG islands and repetitive elements. While methylation at most genomic elements was similar, a striking difference in the distribution of methylation values was observed for first exons and CpG-island promoters between the two strains (**Fig. 1a**). BALB/c mice displayed a notably higher proportion of first exons and CpG-island promoters with few or no methylated bases than C57BL/6 mice. This global effect reflects differences in DNA methylation in approximately 600 genes, which at the level of individual loci can exhibit relatively small differences in promoter and first exon methylation (**Fig. 1b**). The genes in question were not found to be enriched in specific cellular functions or pathways.

**Figure 1.**
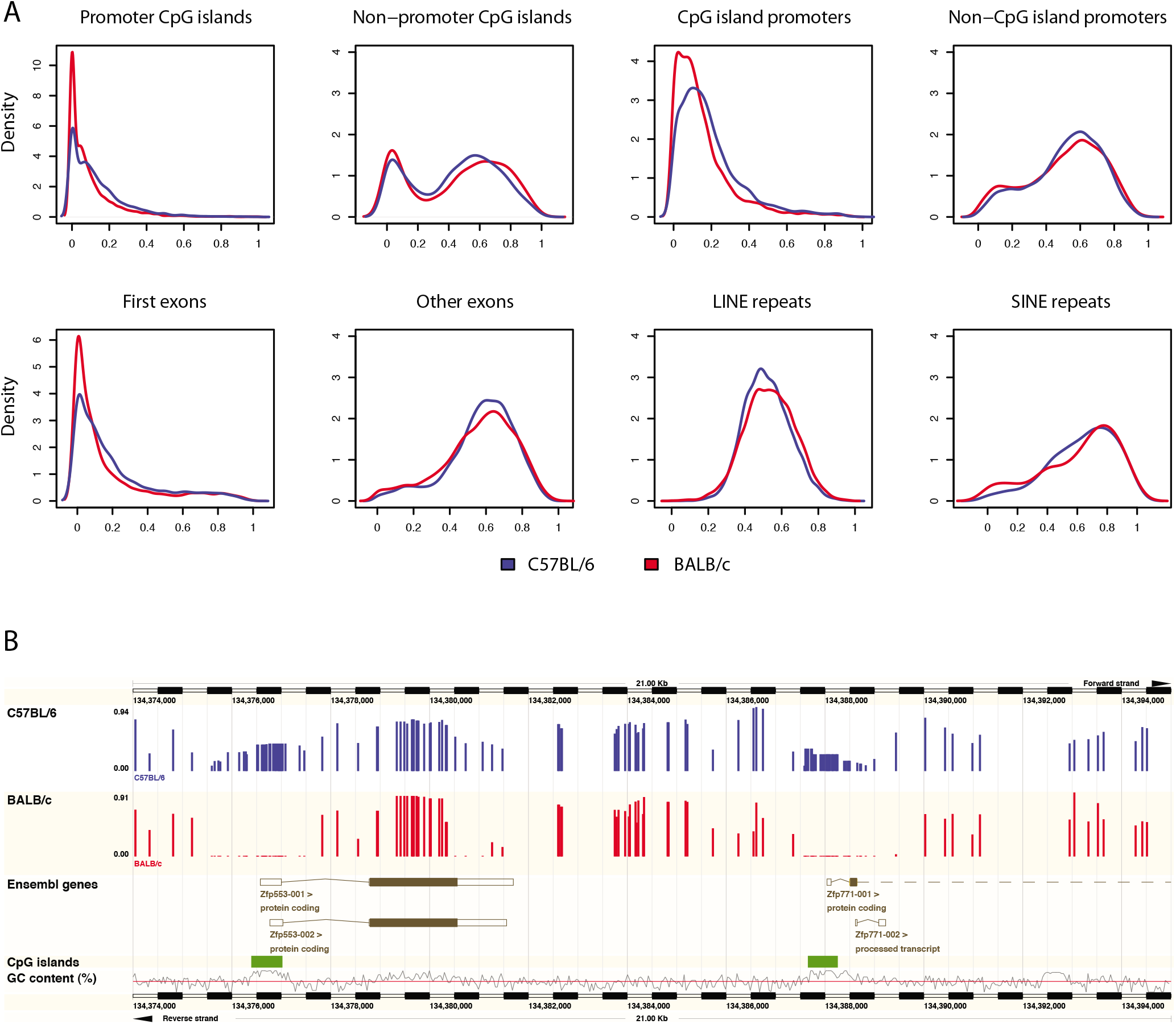
**A**) Distribution of DNA methylation at annotated genomic elements in C57BL/6 and BALB/c mouse brain. **B**) Genome browser image depicting methylation patterns in a region comprising two gene loci. Although methylation levels are similar for most CpGs, higher levels were measured in the promoter regions of C57BL/6 mice.

To assess the extent to which the observed patterns are driven by genetic factors, we performed DNA methylation profiling of offspring from reciprocal crosses between C57BL/6 and BALB/c mice (**Fig. 2a**). Genetic differences at autosomal loci are equalized in the resulting F1 hybrid offspring. Consistent with a genetic contribution to DNA methylation patterning, methylation profiles of C57BL/6×BALB/c and BALB/c×C57BL/6 mice were more similar than those of the pure inbred strains (Pearson r = 0.76, compared to 0.67). As in the inbred strains, about half of all interrogated CpG dinucleotides were found to be partially methylated, although the percentages differed less across the hybrid samples (50.3% in C57BL/6×BALB/c, 53.8% in BALB/c×C57BL/6).

**Figure 2.**
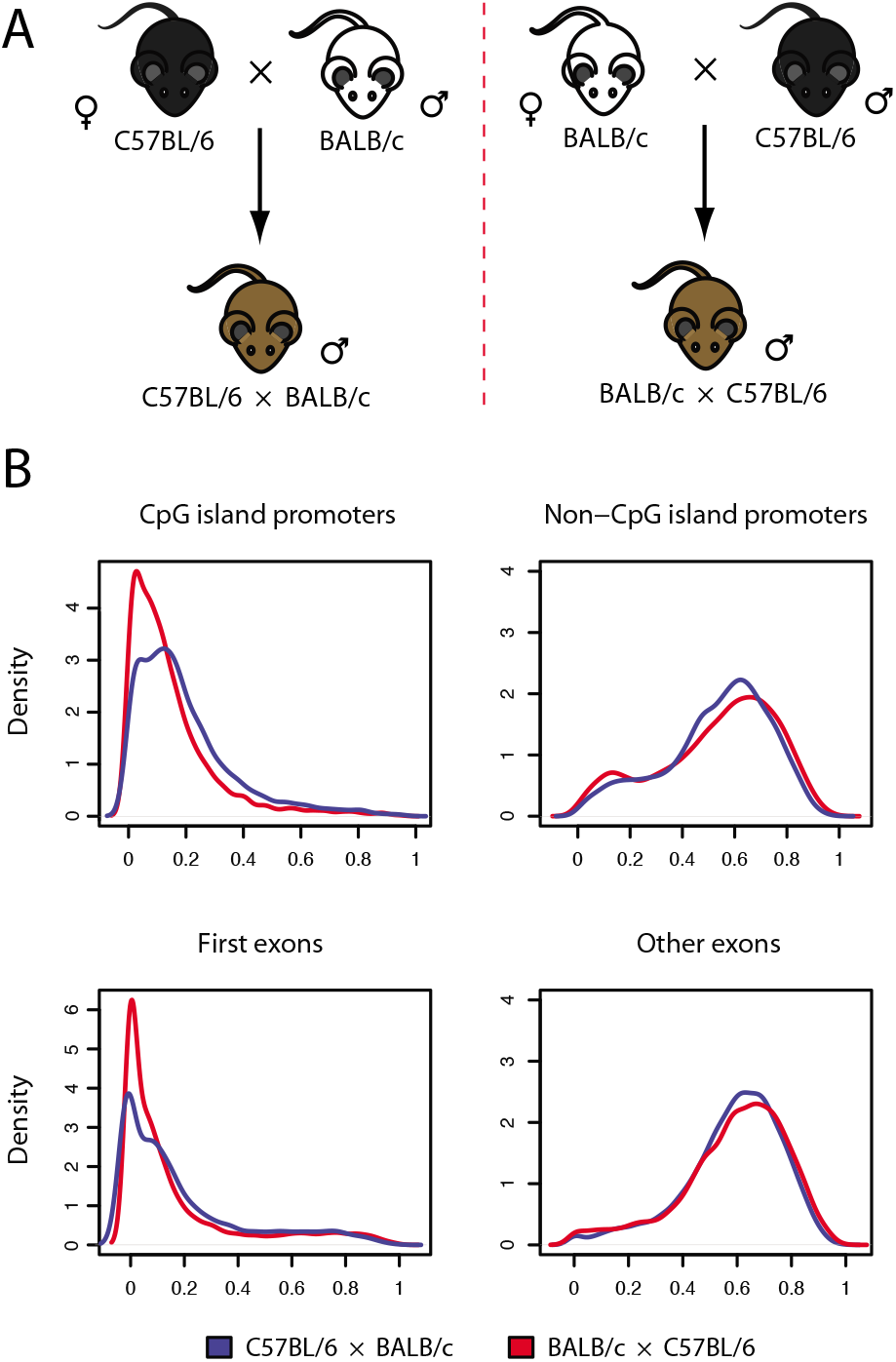
**A**) Experimental design of the comparison between C57BL/6×BALB/c and BALB/c×C57BL/6 reciprocal F1 hybrid mice. **B**) Distribution of DNA methylation at annotated genomic elements in C57BL/6 and BALB/c reciprocal hybrid mouse brain. Hypomethylation of first exons and CpG-island promoters also occurs in F1 hybrids and is determined by the maternal strain.

The greater similarity of the methylation profiles was also reflected in a smaller number (626) of DMRs identified between reciprocal F1 hybrid samples, that could be associated with functional categories involving development and transcriptional regulation (**Supplementary Tables 3,4**).

Similar to what was observed between inbred strains, the distribution of methylation levels indicated a significant increase in first exons and CpG-island promoters with little or no DNA methylation in BALB/c×C57BL/6 when compared to C57BL/6×BALB/c mice (**Fig. 2b**). A comparison of methylation distributions for first exons and CpG islands across all four experimental groups suggested that methylation was correlated with the maternal strain (**Figs. 1a, 2b**). Although male reciprocal F1 hybrid mice differ in the strain of their sex chromosomes, this difference is unlikely to have determined differences in global DNA methylation as these were observed at roughly equal density across the entire genome (**Supplementary Fig. 1**).

Next, we carried out unsupervised hierarchical clustering of DNA methylation profiles to see if evidence for a maternal effect was corroborated at the level of specific genomic elements. Consistent with our finding that overall methylation profiles were primarily driven by genetic differences, clustering based on the methylation level of differentially methylated regions (1000 bp bins) across the entire genome revealed that F1 hybrids were more similar to each other than to either of the inbred strains (**Fig. 3a**). The same trend emerged if the clustering was based on differential methylation across all CpG islands (**Fig. 3b**). However, when performing unsupervised clustering on differentially methylated CpG island promoters, BALB/c×C57BL/6 and pure BALB/C mice are found to be more similar to each other than to any other sample, supporting a potential maternal influence on methylation levels (**Fig. 3c**).

**Figure 3.**
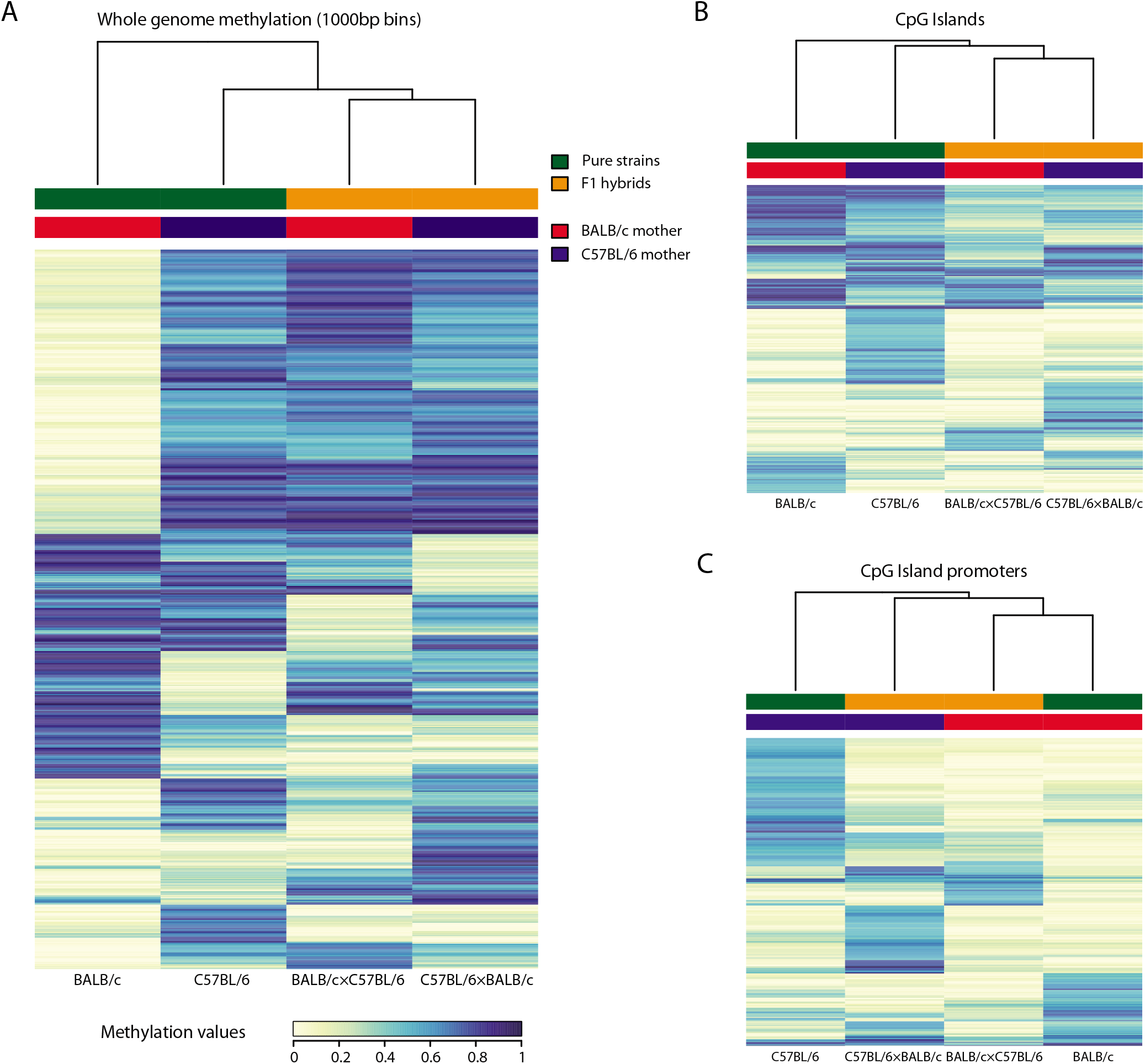
DNA methylation signature of mouse brain tissue in C57BL/6 and BALB/c mice and their F1 hybrids. **A**) Unsupervised hierarchical clustering and heatmap of genome-wide differential methylation, computed over 1000bp regions. Clustering shows the greater similarity of the F1 hybrids. **B)** Unsupervised hierarchical clustering and heatmap of differential CpG island methylation. **C)** Unsupervised hierarchical clustering and heatmap differential methylation of CpG island promoters. While CpG island methylation replicates global trends, CpG island promoters methylation profiles cluster together following the maternal strain.

The association between CpG island promoter methylation profiles and maternal strain could be due to a maternal effect, in which the mother’s non-genetic contribution dictates offspring phenotype. An important source of maternal effects in mammals is the uterine and postnatal maternal environment and C57BL/6 and BALB/c mothers, in particular, are known to show large differences in maternal care that impact the behavior of their offspring. To test whether differences in global methylation detected between reciprocal F1 hybrid mice might be determined by postnatal maternal environment we performed DNA methylation profiling of BALB/c×C57BL/6 F1 hybrid mice cross-fostered after birth to either a C57BL/6 or BALB/c mother (**Fig. 4a**). In these mice, the trend toward global hypomethylation of first exons and CpG island promoter regions was abolished (**Fig. 4b**), demonstrating that postnatal maternal environment differences between strains were not sufficient to reprogram global methylation patterns in offspring and suggesting that uterine or zygotic maternal effects are likely to under the maternal strain-dependent differences we observed.

**Figure 4.**
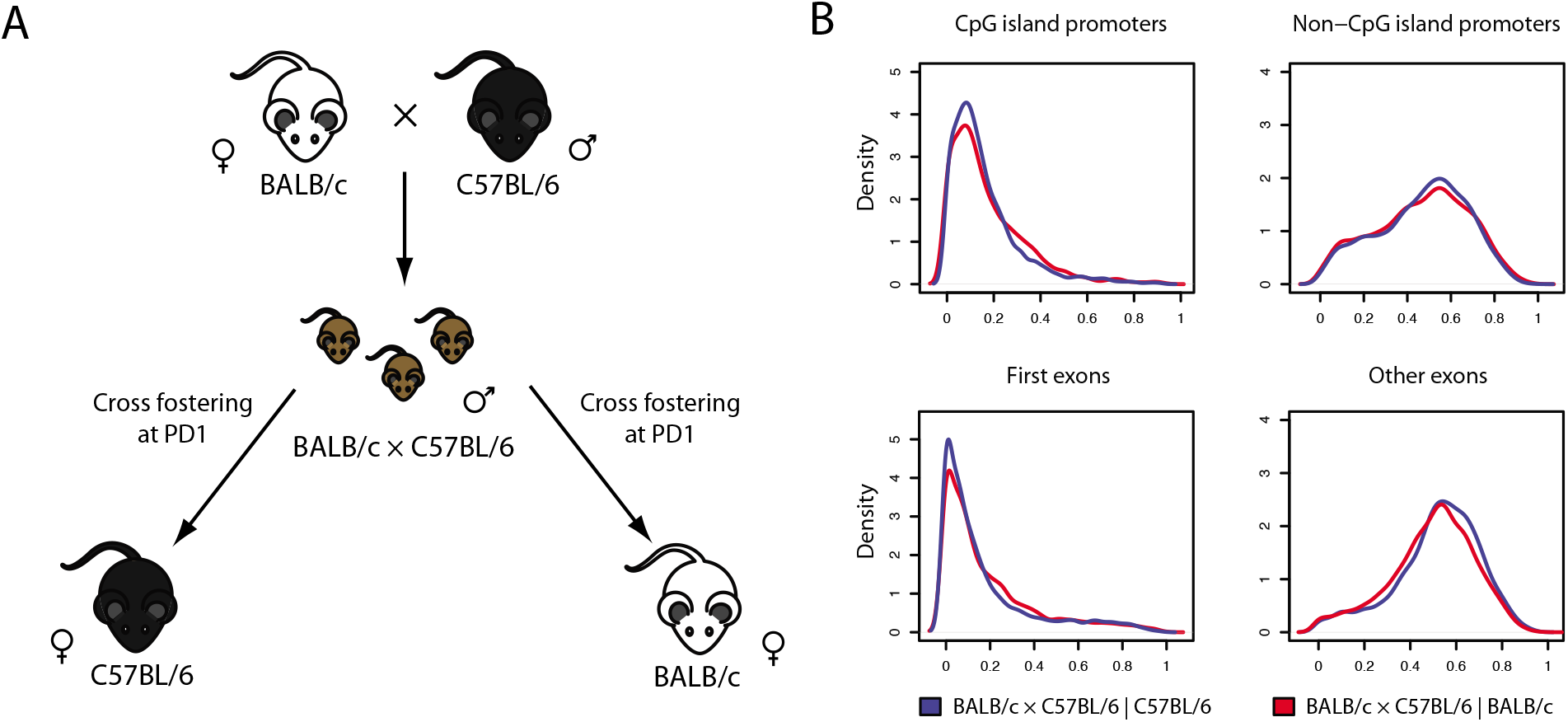
**A**) Experimental design of the comparison between BALB/c×C57BL/6 F1 hybrids cross-fostered after birth to either a C57BL/6 or BALB/c mother. **B**) Distribution of DNA methylation at annotated genomic elements in crossfostered reciprocal hybrid mouse brain. Hypomethylation of first exons and CpG-island promoters is lost in cross-fostered mice.

## Discussion

In this study we aimed to understand the relative impact of genetic and environmental factors on DNA methylation patterns across the mammalian genome in the brain. Inbred strains of mice provide useful models for studying the interaction between genetic and environmental factors in mammals (Carola et al., 2006). We first explored the impact of genetic factors on the global distribution of DNA methylation in C57BL/6 and BALB/c inbred animals. These strains display well-established differences in stress-related behaviors and in susceptibility to the effect of maternal environment on brain development and function (Di Segni et al., 2016). Moreover, C57BL/6 and BALB/c mothers display socially transmitted differences in maternal care (Carola et al., 2006) allowing us to examine potential non-genetic effects on global DNA methylation. Adult hippocampal tissue was chosen for the study as persistent changes in DNA methylation in this structure have been associated with differences in maternal environment (Weaver et al., 2004).

A considerable number of interrogated CpG dinucleotides were found to be in a partially methylated state in both strains. This is consistent with earlier results obtained using whole-genome bisulfite sequencing, which found 40% of all CpGs in fibroblast cells to be partially methylated, although stem cells showed much lower levels (Lister et al., 2009;Lister et al., 2011). Our results suggest that the presence of extensive partially methylated domains could be a common phenomenon in somatic cell types. The higher percentages we find may be partly explained by a greater inherent heterogeneity in tissue samples as compared to cell lines.

Our data demonstrate the existence of significant and stable epigenetic differences between mouse strains, revealed by the >1000 large DMRs that were found throughout the genome between the two strains. These regions are functionally related to embryonic development, chromatin organization, and transcriptional regulation, consistent with the notion that DNA methylation state modulates chromatin accessibility and transcriptional control and plays a crucial role in development. Notably, our findings reveal highly localized DNA methylation differences between the two strains (**Fig. 1b**), with the largest difference in the distribution of methylation observed in first exons and CpG-island promoters (**Fig. 1a**). CpG-island promoters typically show low levels of methylation, a finding that is consistent with their putative evolution as genomic regions protected from cytosine methylation-dependent C-to-T mutation (Edwards et al., 2010). However, DNA methylation at CpG-island promoters is associated with the repression of gene expression and it is thus possible that the differences we report have functional consequences on gene expression.

Consistent with a genetic contribution to DNA methylation patterning (Gertz et al., 2011) methylation profiles of reciprocal F1 offspring were more similar than those of the pure inbred strains (**Fig. 3a,b**). As in pure inbred mice, DMRs could be functionally associated to development and transcriptional regulation, though enrichment in categories related to chromatin organization was no longer observed; this suggests that methylation of chromatin regulators is mainly dependent on genetic background, while regulatory and developmental genes may be more susceptible to epigenetic changes driven by non-genetic factors.

An investigation of the distribution of DNA methylation across specific functional elements in reciprocal F1 hybrid mice, however, revealed a similar pattern of methylation differences as was observed between inbred strains, with methylation levels at CpG-island promoters correlated with the maternal strain. These findings open the possibility of a maternal effect on global DNA methylation patterns in the brain. Maternal effects are phenotypes dictated by the genotype of the mother that are, however, not mediated by her genetic contribution to the zygote. Maternal effects typically involve the deposition of functional gene products by the mother into the zygote or her contribution of the pre- or postnatal environment. In mammals, important maternal effects are generated by the uterine environment and by the maternal care common in these species. Evidence supports the hypothesis that maternal care can effect DNA methylation of genes in both brain and peripheral tissues (Weaver et al., 2004;Arsenault et al., 2018;Lester et al., 2018) and that these effects are associated with persistent changes in gene expression (St-Cyr and McGowan, 2015;Thorsell and Natt, 2016;Elmadih and Abumadini, 2019). However, it remains unclear how specific these changes are and what might be the mechanism of their establishment. Our cross-fostering experiments demonstrated that postnatal maternal environment does not have a major influence on global DNA methylation and suggests that prenatal maternal effects may be responsible for the differences in CpG-island promoter methylation observed between inbred strains. Although our experiments did not allow us to distinguish early zygotic from later uterine maternal effects, zygotic maternal effects have been described in mice that affect global DNA methylation and epigenetic programming, including *Dnmt0* and *Stella* (Han et al., 2019). However, known genetic polymorphisms in these genes between C57BL/6 and BALB/c are not predicted to have functional consequences on gene activity (C. Gross, unpublished data).

In summary, global analysis of DNA methylation patterns in the adult brain revealed that inbred mouse strains harbor significant differences in DNA methylation selectively at first exons and CpG islands of specific classes of genes. These differences were likely to be determined in part by non-genetic maternal factors, as they persist in reciprocal F1 hybrid mice in which autosomal genetic variation was eliminated and correlated with maternal strain. Cross-fostering experiments excluded postnatal environment as a critical maternal factor and pointed to prenatal maternal environment as a potential determinant of global DNA methylation patterns. Further investigations are needed to determine the processes that mediate this effect and their potential impact on gene regulation.

## Acknowledgments

This study was supported by EMBL and NIEHS grant #R21 ES015174. The authors thank Alessandro Guffanti (Genomnia) for technical support, Anne O’Donnell for help in constructing the first Methyl-MAPS libraries, and Tim Bestor for critical advice on the project.

