## supplemental data for "A maternal effect regulates global DNA methylation patterns"

#### Supplemental Figures

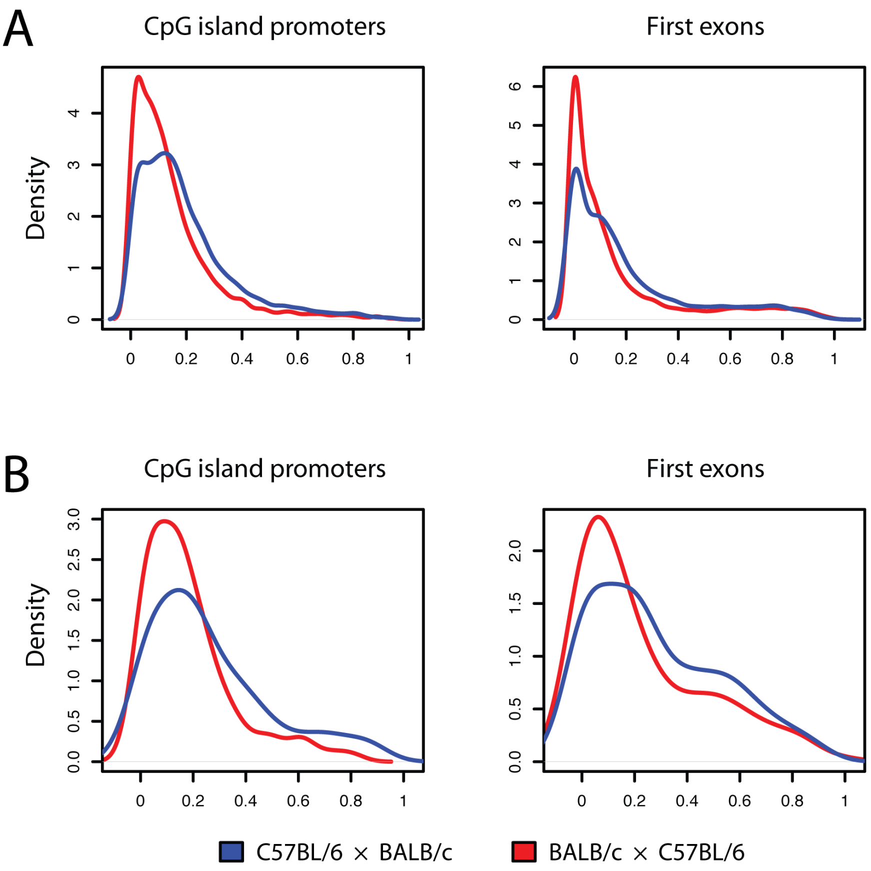


**Supplemental figure 1.** Distributions of methylation values for CpG island promoters and first exons in the C57BL/6×BALB/c - BALB/c×C57BL/6 comparison, considering (A) only autosomes and (B) only sex chromosomes.

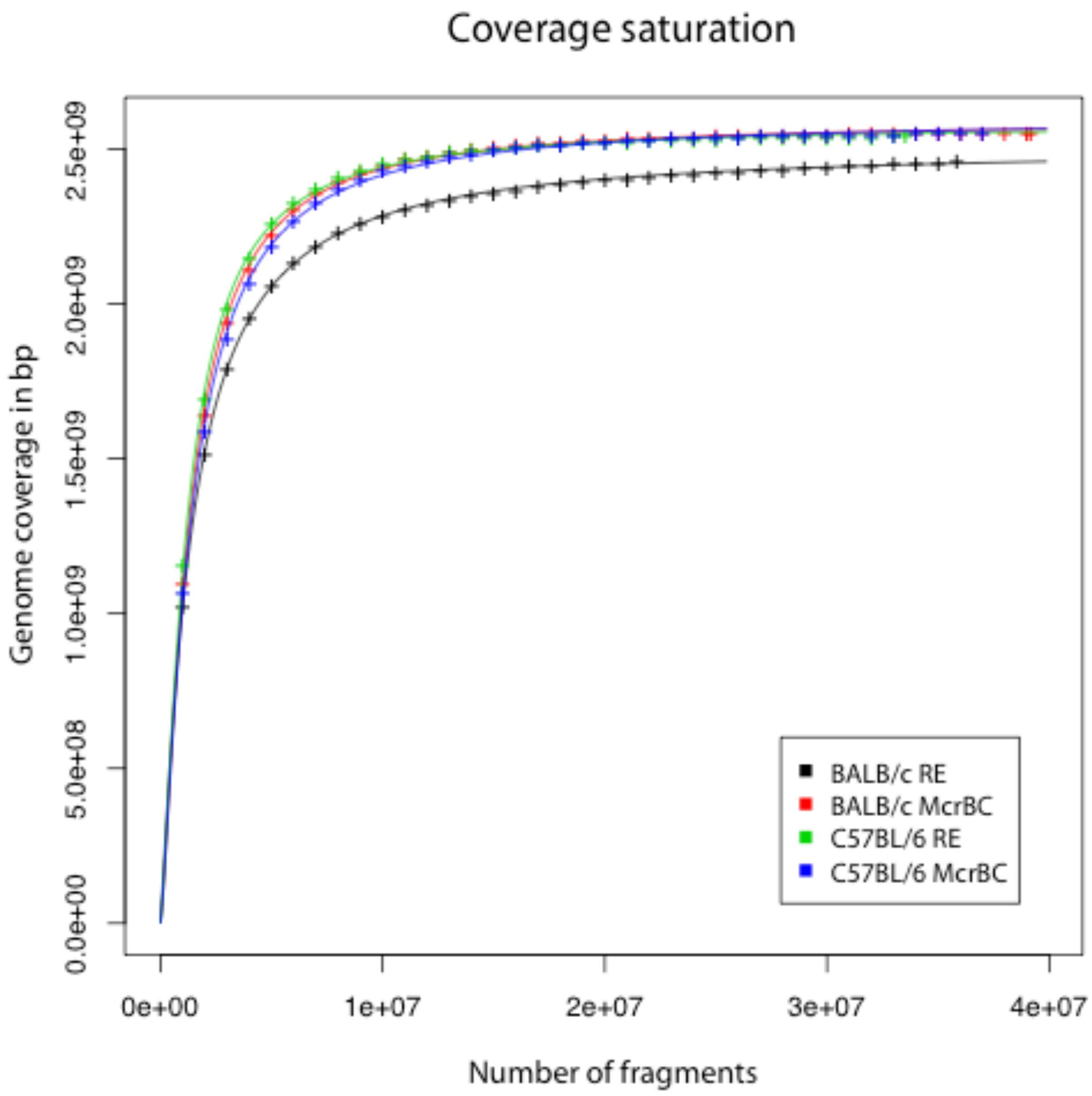


**Supplemental figure 2.** Genome coverage of samples of the RE and McrBC fragments of the C57BL/6 - BALB/c comparison, fitted by a saturation curve. Half of the fragments provide >98% of the coverage, showing that the genome coverage computed from the data in this study is a good approximation of the actual expected coverage across the genome.

### Supplemental Tables

Due to their size, supplemental tables 1 and 3 are provided in separate files.

**Supplemental table 1**. Genes containing a differentially methylated region either in the gene body or within a defined promoter-proximal region (< 5Kb upstream from TSS) in BALB/c - C57BL/6 comparison.

**Supplemental table 2**. Gene Ontology (GO) analysis for genes containing a differentially methylated region in the gene body or promoter region in BALB/c - C57BL/6 comparison.

| **Functional term** | **Genes** | **Percentage** | **P-value** | **Fold enrichment** | **Adjusted P-value** (Benjamini-Hochberg) |
|---|---|---|---|---|---|
| GO:0009987~cellular process | 281 | 51.09 | 7.28E-11 | 1.24 | 1.31E-07 |
| GO:0048598~embryonic morphogenesis | 33 | 6 | 2.42E-10 | 3.75 | 2.18E-07 |
| GO:0048568~embryonic organ development | 26 | 4.72 | 1.08E-09 | 4.40 | 6.52E-07 |
| GO:0019222~regulation of metabolic process | 120 | 21.81 | 1.78E-09 | 1.66 | 8.03E-07 |
| GO:0048856~anatomical structure development | 95 | 17.27 | 7.07E-09 | 1.78 | 2.55E-06 |
| GO:0048731~system development | 90 | 16.36 | 1.09E-08 | 1.81 | 3.28E-06 |
| GO:0048513~organ development | 78 | 14.18 | 1.53E-08 | 1.91 | 3.95E-06 |
| GO:0080090~regulation of primary metabolic process | 109 | 19.81 | 1.61E-08 | 1.66 | 3.64E-06 |
| GO:0009887~organ morphogenesis | 40 | 7.27 | 1.77E-08 | 2.75 | 3.56E-06 |
| GO:0031323~regulation of cellular metabolic process | 112 | 20.36 | 2.05E-08 | 1.64 | 3.71E-06 |
| GO:0060255~regulation of macromolecule metabolic process | 109 | 19.81 | 2.25E-08 | 1.65 | 3.68E-06 |
| GO:0009790~embryonic development | 42 | 7.63 | 5.32E-08 | 2.56 | 7.99E-06 |
| GO:0006333~chromatin assembly or disassembly | 16 | 2.90 | 6.10E-08 | 5.99 | 8.46E-06 |
| GO:0048562~embryonic organ morphogenesis | 19 | 3.45 | 7.84E-08 | 4.82 | 1.01E-05 |
| GO:0045449~regulation of transcription | 93 | 16.90 | 1.09E-07 | 1.70 | 1.32E-05 |
| GO:0010468~regulation of gene expression | 99 | 18 | 1.13E-07 | 1.66 | 1.28E-05 |
| GO:0007389~pattern specification process | 25 | 4.54 | 1.27E-07 | 3.59 | 1.35E-05 |
| GO:0009653~anatomical structure morphogenesis | 57 | 10.36 | 1.52E-07 | 2.09 | 1.52E-05 |
| GO:0010556~regulation of macromolecule biosynthetic process | 97 | 17.63 | 1.67E-07 | 1.66 | 1.59E-05 |
| GO:0007275~multicellular organismal development | 101 | 18.36 | 1.84E-07 | 1.63 | 1.66E-05 |
| GO:0019219~regulation of nucleobase, nucleoside, nucleotide and nucleic acid metabolic process | 96 | 17.45 | 1.88E-07 | 1.66 | 1.61E-05 |
| GO:0051252~regulation of RNA metabolic process | 69 | 12.54 | 2.21E-07 | 1.89 | 1.81E-05 |
| GO:0006355~regulation of transcription, DNA-dependent | 68 | 12.36 | 2.70E-07 | 1.89 | 2.12E-05 |

| | | | | | |
|---|---|---|---|---|---|
| GO:0051171~regulation of nitrogen compound metabolic process | 96 | 17.45 | 2.75E-07 | 1.65 | 2.06E-05 |
| GO:0031326~regulation of cellular biosynthetic process | 98 | 17.81 | 4.68E-07 | 1.62 | 3.37E-05 |
| GO:0009889~regulation of biosynthetic process | 98 | 17.81 | 5.55E-07 | 1.61 | 3.85E-05 |
| GO:0006334~nucleosome assembly | 12 | 2.181 | 1.44E-06 | 6.71 | 9.59E-05 |
| GO:0032502~developmental process | 104 | 18.90 | 1.65E-06 | 1.54 | 1.06E-04 |
| GO:0031497~chromatin assembly | 12 | 2.18 | 1.89E-06 | 6.53 | 1.18E-04 |
| GO:0065004~protein-DNA complex assembly | 12 | 2.18 | 2.17E-06 | 6.45 | 1.30E-04 |
| GO:0034728~nucleosome organization | 12 | 2.18 | 2.17E-06 | 6.45 | 1.30E-04 |
| GO:0010467~gene expression | 97 | 17.66 | 5.63E-06 | 1.54 | 3.28E-04 |
| GO:0006996~organelle organization | 51 | 9.27 | 7.73E-06 | 1.93 | 4.36E-04 |
| GO:0044260~cellular macromolecule metabolic process | 150 | 27.27 | 9.71E-06 | 1.34 | 5.31E-04 |
| GO:0006325~chromatin organization | 23 | 4.18 | 9.95E-06 | 2.98 | 5.28E-04 |
| GO:0001822~kidney development | 13 | 2.36 | 1.14E-05 | 4.96 | 5.89E-04 |
| GO:0048522~positive regulation of cellular process | 58 | 10.54 | 1.43E-05 | 1.79 | 7.18E-04 |
| GO:0003002~regionalization | 18 | 3.27 | 2.02E-05 | 3.43 | 9.84E-04 |
| GO:0006350~transcription | 71 | 12.90 | 2.61E-05 | 1.63 | 0.001239435 |
| GO:0045595~regulation of cell differentiation | 25 | 4.54 | 2.73E-05 | 2.64 | 0.001260688 |
| GO:0043170~macromolecule metabolic process | 162 | 29.45 | 3.26E-05 | 1.29 | 0.001467294 |
| GO:0006323~DNA packaging | 12 | 2.18 | 3.49E-05 | 4.85 | 0.001534655 |
| GO:0009888~tissue development | 34 | 6.18 | 3.91E-05 | 2.17 | 0.001677301 |
| GO:0043009~chordate embryonic development | 26 | 4.72 | 3.97E-05 | 2.52 | 0.00166414 |
| GO:0009792~embryonic development ending in birth or egg hatching | 26 | 4.72 | 4.65E-05 | 2.49 | 0.001906233 |
| GO:0034645~cellular macromolecule biosynthetic process | 88 | 16 | 5.50E-05 | 1.49 | 0.002200747 |
| GO:0051276~chromosome organization | 25 | 4.54 | 5.68E-05 | 2.52 | 0.002226666 |
| GO:0009059~macromolecule biosynthetic process | 88 | 16 | 6.21E-05 | 1.49 | 0.002381685 |
| GO:0016043~cellular component organization | 73 | 13.27 | 6.52E-05 | 1.57 | 0.002446033 |
| GO:0009952~anterior/posterior pattern formation | 14 | 2.54 | 9.63E-05 | 3.73 | 0.003539825 |
| GO:0006357~regulation of transcription from RNA polymerase II promoter | 32 | 5.81 | 1.13E-04 | 2.12 | 0.00406362 |
| GO:0007399~nervous system development | 40 | 7.27 | 1.17E-04 | 1.91 | 0.004122026 |
| GO:0060429~epithelium development | 19 | 3.45 | 1.22E-04 | 2.86 | 0.004214266 |
| GO:0050793~regulation of developmental process | 30 | 5.45 | 1.45E-04 | 2.15 | 0.004938889 |
| GO:0060541~respiratory system development | 12 | 2.18 | 2.26E-04 | 3.95 | 0.007512887 |
| GO:0048518~positive regulation of biological process | 59 | 10.72 | 2.29E-04 | 1.61 | 0.007472267 |

| | | | | | |
|---|---|---|---|---|---|
| GO:0001655~urogenital system development | 13 | 2.36 | 2.43E-04 | 3.63 | 0.007788413 |
| GO:0030154~cell differentiation | 61 | 11.09 | 2.64E-04 | 1.58 | 0.008333204 |
| GO:0048869~cellular developmental process | 63 | 11.45 | 2.66E-04 | 1.57 | 0.008236027 |
| GO:0035295~tube development | 18 | 3.27 | 2.70E-04 | 2.78 | 0.008231772 |
| GO:0034622~cellular macromolecular complex assembly | 16 | 2.90 | 2.89E-04 | 3.01 | 0.008663242 |
| GO:0002009~morphogenesis of an epithelium | 14 | 2.54 | 3.27E-04 | 3.30 | 0.009631827 |
| GO:0043010~camera-type eye development | 12 | 2.18 | 3.41E-04 | 3.77 | 0.009866919 |
| GO:0034621~cellular macromolecular complex subunit organization | 17 | 3.09 | 3.48E-04 | 2.83 | 0.009905135 |

**Supplemental table 3**. Genes containing a differentially methylated region either in the gene body or within a defined promoter-proximal region (< 5Kb upstream from TSS) in the BALB/c×C57BL/6 -C57BL/6×BALB/c comparison.

**Supplemental table 4**. Gene Ontology (GO) analysis for genes with a differentially methylated region in the gene body or promoter region in the BALB/c×C57BL/6 -C57BL/6×BALB/c comparison.

| **Functional term** | **Genes** | **Percentage** | **P-value** | **Fold enrichment** | **Adjusted P-value** (Benjamini-Hochberg) |
|---|---|---|---|---|---|
| GO:0006355~regulation of transcription, DNA-dependent | 41 | 14.96 | 2.02E-06 | 2.19 | 0.00269333 |
| GO:0051252~regulation of RNA metabolic process | 41 | 14.96 | 2.97E-06 | 2.16 | 0.001985972 |
| GO:0007417~central nervous system development | 18 | 6.56 | 4.55E-06 | 3.83 | 0.002026146 |
| GO:0045449~regulation of transcription | 53 | 19.34 | 4.56E-06 | 1.86 | 0.001522201 |
| GO:0019219~regulation of nucleobase, nucleoside, nucleotide and nucleic acid metabolic process | 55 | 20.07 | 4.66E-06 | 1.83 | 0.001245552 |
| GO:0051171~regulation of nitrogen compound metabolic process | 55 | 20.07 | 5.95E-06 | 1.82 | 0.001324477 |
| GO:0048731~system development | 49 | 17.88 | 8.37E-06 | 1.89 | 0.001598124 |
| GO:0009653~anatomical structure morphogenesis | 33 | 12.04 | 9.45E-06 | 2.32 | 0.001577891 |
| GO:0007399~nervous system development | 28 | 10.21 | 9.58E-06 | 2.57 | 0.001422672 |
| GO:0007420~brain development | 15 | 5.47 | 1.80E-05 | 4.09 | 0.002408819 |
| GO:0048856~anatomical structure development | 50 | 18.24 | 2.39E-05 | 1.81 | 0.002905076 |
| GO:0010468~regulation of gene expression | 54 | 19.70 | 2.58E-05 | 1.74 | 0.00286876 |
| GO:0010556~regulation of macromolecule biosynthetic process | 53 | 19.34 | 3.10E-05 | 1.74 | 0.003184623 |
| GO:0031326~regulation of cellular biosynthetic process | 54 | 19.70 | 3.99E-05 | 1.71 | 0.003800675 |
| GO:0048513~organ development | 41 | 14.96 | 4.17E-05 | 1.93 | 0.003713343 |
| GO:0009889~regulation of biosynthetic process | 54 | 19.70 | 4.41E-05 | 1.71 | 0.003679986 |
| GO:0030154~cell differentiation | 39 | 14.23 | 5.88E-05 | 1.95 | 0.004610336 |

| | | | | | |
|---|---|---|---|---|---|
| GO:0007275~multicellular organismal development | 54 | 19.70 | 7.23E-05 | 1.68 | 0.005359277 |
| GO:0031323~regulation of cellular metabolic process | 58 | 21.16 | 7.42E-05 | 1.63 | 0.005206343 |
| GO:0032502~developmental process | 57 | 20.80 | 9.41E-05 | 1.63 | 0.006272762 |
| GO:0048869~cellular developmental process | 39 | 14.23 | 1.43E-04 | 1.87 | 0.0090369 |
| GO:0006807~nitrogen compound metabolic process | 63 | 22.99 | 1.52E-04 | 1.54 | 0.009171382 |
| GO:0030182~neuron differentiation | 16 | 5.83 | 1.70E-04 | 3.15 | 0.009826177 |
| GO:0080090~regulation of primary metabolic process | 55 | 20.07 | 1.75E-04 | 1.61 | 0.009690132 |
| GO:0019222~regulation of metabolic process | 59 | 21.53 | 1.79E-04 | 1.57 | 0.009523851 |
| GO:0009887~organ morphogenesis | 20 | 7.29 | 1.92E-04 | 2.64 | 0.009841653 |

**Supplemental table 5.** Numbers of reads, identified fragments, and interrogated CpGs for each sample. Arrows indicate cross-fostering experiments.

| Sample | Sequencing reads | Filtered fragments | CpGs |
|---|---|---|---|
| C57BL/6 RE | 159,633,174 (2×79.8 M) | 33,517,898 | 5,486,645 |
| C57BL/6 McrBC | 160,909,598 (2×80.5 M) | 36,978,783 | |
| BALB/c RE | 165,229,480 (2×82.6 M) | 35,849,523 | 5,487,076 |
| BALB/c McrBC | 165,390,658 (2×82.7 M) | 39,172,297 | |
| C57BL/6×BALB/c RE | 129,255,064 (2×64.6 M) | 32,047,946 | 5,382,490 |
| C57BL/6×BALB/c McrBC | 116,735,676 (2×58.4 M) | 21,409,398 | |
| BALB/c×C57BL/6 RE | 110,404,456 (2×55.2 M) | 27,366,315 | 5,460,661 |
| BALB/c×C57BL/6 McrBC | 104,517,490 (2×52.3 M) | 20,740,679 | |
| BALB/c×C57BL/6 → C57BL/6 RE | 171,119,945 (2×85.6 M) | 31,535,898 | 5,530,783 |
| BALB/c×C57BL/6 → C57BL/6 McrBC | 174,159,099 (2×87.1 M) | 32,978,239 | |
| BALB/c×C57BL/6 → BALB/c RE | 177,101,623 (2×88.6 M) | 26,746,965 | 5,511,126 |
| BALB/c×C57BL/6 → BALB/c McrBC | 172,939,393 (2×86.5 M) | 28,275,441 | |
