## supplementary table 1 for "A maternal effect regulates global DNA methylation patterns"

**Table 1a**. List of genes containing a hypomethylated region in gene body or promoter region (< 5Kb upstream from TSS) in BALB/c compared to C57BL/6.

| EnsEmbl ID | Gene name | Entrez ID |
|---|---|---|
| ENSMUSG00000059975 | Zfp74 | 72723 |
| ENSMUSG00000030643 | Rab30 | 75985 |
| ENSMUSG00000066721 | Zfp575 | 101544 |
| ENSMUSG00000059273 | Zc3h4 | 330474 |
| ENSMUSG00000030657 | Xylt1 | 233781 |
| ENSMUSG00000055026 | Gabrg3 | 14407 |
| ENSMUSG00000078801 | Gm10991 | |
| ENSMUSG00000030507 | Dbx1 | 13172 |
| ENSMUSG00000091898 | Tnnc1 | 21924 |
| ENSMUSG00000039197 | Adk | 11534 |
| ENSMUSG00000021824 | Ap3m1 | 55946 |
| ENSMUSG00000090351 | Gm17204 | |
| ENSMUSG00000022021 | Diap3 | 56419 |
| ENSMUSG00000035566 | Pcdh17 | 219228 |
| ENSMUSG00000084925 | 1810062O18Rik | |
| ENSMUSG00000084276 | Gm8317 | |
| ENSMUSG00000022021 | Diap3 | 56419 |
| ENSMUSG00000022113 | Trim52 | 212085 |
| ENSMUSG00000062093 | Gm10110 | 100503296 |
| ENSMUSG00000047171 | Helt | 234219 |
| ENSMUSG00000055932 | Fto | 26383 |
| ENSMUSG00000089704 | Galnt2 | 108148 |
| ENSMUSG00000086717 | Gm15655 | |
| ENSMUSG00000055435 | Maf | 17132 |
| ENSMUSG00000055932 | Fto | 26383 |
| ENSMUSG00000036356 | Csgalnact1 | 234356 |
| ENSMUSG00000047495 | Dlgap2 | 244310 |
| ENSMUSG00000071644 | Eef1g | 67160 |
| ENSMUSG00000055491 | Pprc1 | 226169 |
| ENSMUSG00000044220 | Nkx2-3 | 18089 |
| ENSMUSG00000024985 | Tcf7l2 | 21416 |
| ENSMUSG00000056612 | Ppp1r14b | 18938 |
| ENSMUSG00000067608 | Pcna-ps2 | |
| ENSMUSG00000084364 | Gm15801 | |
| ENSMUSG00000025007 | Aldh18a1 | 56454 |
| ENSMUSG00000026064 | Ptp4a1 | 19243 |
| ENSMUSG00000040323 | Gm15429 | |
| ENSMUSG00000054203 | Ifi205 | 226695 |
| ENSMUSG00000033774 | Npbwr1 | 226304 |
| ENSMUSG00000025983 | Ccdc150 | 78016 |
| ENSMUSG00000038034 | Igsf8 | 140559 |
| ENSMUSG00000043716 | Rpl7 | 19989 |
| ENSMUSG00000025921 | Rdh10 | 98711 |
| ENSMUSG00000060743 | H3f3a | 15081 |
| ENSMUSG00000026039 | Sgol2 | 68549 |
| ENSMUSG00000041377 | Ninj2 | 29862 |
| ENSMUSG00000071477 | Zfp777 | 72306 |
| ENSMUSG00000068545 | Gm10243 | |
| ENSMUSG00000044211 | Gm7887 | |

| | | |
|---|---|---|
| ENSMUSG00000030007 | Cct7 | 12468 |
| ENSMUSG00000030008 | 1700040I03Rik | 73327 |
| ENSMUSG00000073067 | 9130019P16Rik | |
| ENSMUSG00000087380 | 2210408F21Rik | |
| ENSMUSG00000000093 | Tbx2 | 21385 |
| ENSMUSG00000040850 | Psme4 | 103554 |
| ENSMUSG00000044795 | Cyb5d1 | 327951 |
| ENSMUSG00000018476 | Kdm6b | 216850 |
| ENSMUSG00000045377 | Tmem88 | 67020 |
| ENSMUSG00000059278 | Lsmd1 | 78304 |
| ENSMUSG00000013415 | Igf2bp1 | 140486 |
| ENSMUSG00000020160 | Meis1 | 17268 |
| ENSMUSG00000041695 | Kcnj2 | 16518 |
| ENSMUSG00000000093 | Tbx2 | 21385 |
| ENSMUSG00000065529 | Mir22 | |
| ENSMUSG00000038217 | Tlcd2 | 380712 |
| ENSMUSG00000085148 | 2210403K04Rik | |
| ENSMUSG00000044068 | Zrsr1 | 22183 |
| ENSMUSG00000024112 | Cacna1h | 58226 |
| ENSMUSG00000086895 | 9330136K24Rik | |
| ENSMUSG00000024114 | Prss41 | 71003 |
| ENSMUSG00000048992 | Prss32 | 69814 |
| ENSMUSG00000071256 | Zfp213 | 449521 |
| ENSMUSG00000090725 | Gm17633 | |
| ENSMUSG00000079594 | Gm1975 | 100038854 |
| ENSMUSG00000088517 | U6 | |
| ENSMUSG00000022964 | Tmem50b | 77975 |
| ENSMUSG00000022706 | Mrpl40 | 18100 |
| ENSMUSG00000022702 | Hira | 15260 |
| ENSMUSG00000073563 | Csnk1g3 | 70425 |
| ENSMUSG00000023036 | Pcdhga11 | 93724 |
| ENSMUSG00000074513 | Arfip1 | 99889 |
| ENSMUSG00000027833 | Shox2 | 20429 |
| ENSMUSG00000056947 | Mab21l1 | 17116 |
| ENSMUSG00000087825 | U6 | |
| ENSMUSG00000074580 | 4931440P22Rik | |
| ENSMUSG00000034640 | Tiparp | 99929 |
| ENSMUSG00000056260 | 4933421E11Rik | 321000 |
| ENSMUSG00000078684 | 5830417I10Rik | |
| ENSMUSG00000074513 | Arfip1 | 99889 |
| ENSMUSG00000028044 | Cks1b | 54124 |
| ENSMUSG00000042626 | Shc1 | 20416 |
| ENSMUSG00000087440 | 4930577N17Rik | |
| ENSMUSG00000050936 | Hist2h2bb | |
| ENSMUSG00000074403 | Hist2h3b | 319154 |
| ENSMUSG00000027833 | Shox2 | 20429 |
| ENSMUSG00000015750 | Aph1a | 226548 |
| ENSMUSG00000061911 | Myt1l | 17933 |
| ENSMUSG00000087109 | 4930474H06Rik | |
| ENSMUSG00000059229 | Gm6802 | |
| ENSMUSG00000085622 | 3110056K07Rik | |
| ENSMUSG00000048118 | Arid4a | 238247 |
| ENSMUSG00000052673 | Gm9887 | |

| | | |
|---|---|---|
| ENSMUSG00000044147 | Arf6 | 11845 |
| ENSMUSG00000020594 | Pum2 | 80913 |
| ENSMUSG00000003352 | Cacnb3 | 12297 |
| ENSMUSG00000060992 | Copz1 | 56447 |
| ENSMUSG00000037386 | Rims2 | 116838 |
| ENSMUSG00000023022 | Lima1 | 65970 |
| ENSMUSG00000079641 | Rpl39 | 100504784 |
| ENSMUSG00000065642 | Snora69 | |
| ENSMUSG00000083411 | Rpl30-ps10 | |
| ENSMUSG00000080968 | Gm1862 | |
| ENSMUSG00000042271 | Nxt2 | 237082 |
| ENSMUSG00000083994 | Gm16465 | |
| ENSMUSG00000079704 | Gm14459 | 546263 |
| ENSMUSG00000064398 | U2 | |
| ENSMUSG00000031060 | Rbm10 | 433287 |
| ENSMUSG00000031059 | Ndufb11 | 104130 |
| ENSMUSG00000081477 | Gm7916 | |
| ENSMUSG00000038344 | Txlng | 353170 |
| ENSMUSG00000000037 | Scml2 | 107815 |
| ENSMUSG00000083130 | Gm14560 | |
| ENSMUSG00000081159 | Gm6023 | |
| ENSMUSG00000031262 | Cenpi | 102920 |
| ENSMUSG00000045103 | Dmd | 13405 |
| ENSMUSG00000073889 | Il11ra1 | 16157 |
| ENSMUSG00000028532 | Cachd1 | 320508 |
| ENSMUSG00000083724 | Gm12651 | |
| ENSMUSG00000085037 | 4933421O10Rik | |
| ENSMUSG00000028277 | Ube2j1 | 56228 |
| ENSMUSG00000028717 | Tal1 | 21349 |
| ENSMUSG00000051351 | Zfp46 | 22704 |
| ENSMUSG00000028782 | Bai2 | 230775 |
| ENSMUSG00000086096 | Gm12688 | |
| ENSMUSG00000027296 | Itpka | 228550 |
| ENSMUSG00000064722 | SNORA44 | |
| ENSMUSG00000065511 | Mir129-2 | |
| ENSMUSG00000027423 | Snx5 | 69178 |
| ENSMUSG00000027424 | 8430406I07Rik | 74528 |
| ENSMUSG00000077714 | Snord17 | |
| ENSMUSG00000033705 | Stard9 | |
| ENSMUSG00000070692 | Gm14059 | |
| ENSMUSG00000079966 | Gm14058 | |
| ENSMUSG00000090313 | Gm17325 | |
| ENSMUSG00000034701 | Neurod1 | 18012 |
| ENSMUSG00000037143 | 4930529M08Rik | 100503181 |
| ENSMUSG00000081406 | Gm13654 | |
| ENSMUSG00000082056 | Gm13745 | |
| ENSMUSG00000045225 | Olfr1152 | 258103 |
| ENSMUSG00000082056 | Gm13745 | |
| ENSMUSG00000026748 | Plxdc2 | 67448 |
| ENSMUSG00000027523 | Gnas | 14683 |
| ENSMUSG00000045994 | B3gat1 | 76898 |
| ENSMUSG00000032011 | Thy1 | 21838 |
| ENSMUSG00000037742 | Eef1a1 | 13627 |

| | | |
|---|---|---|
| ENSMUSG00000076138 | mmu-mir-703 | |
| ENSMUSG00000090470 | 2410012M07Rik | 71979 |
| ENSMUSG00000066477 | Gm16551 | |
| ENSMUSG00000025240 | Sacm1l | 83493 |
| ENSMUSG00000043013 | Onecut1 | 15379 |
| ENSMUSG00000084663 | Mir1900 | |
| ENSMUSG00000032174 | Icam5 | 15898 |
| ENSMUSG00000001014 | Icam4 | 78369 |
| ENSMUSG00000091118 | Mir1900 | |
| ENSMUSG00000059288 | Cdyl | 12593 |
| ENSMUSG00000069274 | Hist1h4f | 100041230 |
| ENSMUSG00000052565 | Hist1h1d | 14957 |
| ENSMUSG00000062808 | Hist1h3d | 319154 |
| ENSMUSG00000069272 | Hist1h2ae | 665433 |
| ENSMUSG00000069268 | Hist1h2bf | 319187 |
| ENSMUSG00000058385 | Hist1h2bg | 319189 |
| ENSMUSG00000071478 | Hist1h2ad | 665433 |
| ENSMUSG00000025408 | Ddit3 | 13198 |
| ENSMUSG00000058799 | Nap1l1 | 53605 |
| ENSMUSG00000045680 | Tcf21 | 21412 |
| ENSMUSG00000020014 | 4930485B16Rik | 380654 |
| ENSMUSG00000025353 | Ormdl2 | 433874 |
| ENSMUSG00000078427 | Sarnp | 66118 |
| ENSMUSG00000079463 | Gm17682 | |
| ENSMUSG00000029723 | Tsc22d4 | 78829 |
| ENSMUSG00000029385 | Ccng2 | 12452 |
| ENSMUSG00000032959 | Pebp1 | 23980 |
| ENSMUSG00000039095 | En2 | 13799 |

**Table 1b**. List of genes containing a hypermethylated region in gene body or promoter region (< 5Kb upstream from TSS) in BALB/c compared to C57BL/6.

| EnsEmbl ID | Gene name | Entrez ID |
|---|---|---|
| ENSMUSG00000049643 | 2310022A10Rik | 66367 |
| ENSMUSG00000040811 | Eml2 | 72205 |
| ENSMUSG00000008028 | 1700008O03Rik | 69349 |
| ENSMUSG00000035064 | Eef2k | 13631 |
| ENSMUSG00000046591 | 5730590G19Rik | 77011 |
| ENSMUSG00000038502 | Ptov1 | 84113 |
| ENSMUSG00000066306 | Numa1 | 101706 |
| ENSMUSG00000007837 | Prrg2 | 65116 |
| ENSMUSG00000073858 | Itpripl2 | 319622 |
| ENSMUSG00000073859 | Itpripl2 | |
| ENSMUSG00000048078 | Odz4 | 23966 |
| ENSMUSG00000089230 | U7 | |
| ENSMUSG00000054676 | 1600014C10Rik | 72244 |
| ENSMUSG00000040811 | Eml2 | 72205 |
| ENSMUSG00000045039 | Megf8 | 269878 |
| ENSMUSG00000033917 | Gde1 | 56209 |

| | | |
|---|---|---|
| ENSMUSG00000064231 | Gm5321 | 384525 |
| ENSMUSG00000036570 | Fxyd1 | 56188 |
| ENSMUSG00000006311 | Etv2 | 14008 |
| ENSMUSG00000038738 | Shank1 | 243961 |
| ENSMUSG00000043671 | Dpy19l3 | 233115 |
| ENSMUSG00000030551 | Nr2f2 | 11819 |
| ENSMUSG00000003273 | Car11 | 12348 |
| ENSMUSG00000059824 | Dbp | 13170 |
| ENSMUSG00000047085 | Lrrc4b | 272381 |
| ENSMUSG00000036291 | Mudeng | 74385 |
| ENSMUSG00000022100 | Xpo7 | 65246 |
| ENSMUSG00000042104 | Uggt2 | 66435 |
| ENSMUSG00000035960 | Apex1 | 11792 |
| ENSMUSG00000007817 | Zmiz1 | 328365 |
| ENSMUSG00000090779 | Gm17110 | |
| ENSMUSG00000021945 | Zmym2 | 76007 |
| ENSMUSG00000021840 | Mapk1ip1l | 218975 |
| ENSMUSG00000039081 | Zfp503 | 218820 |
| ENSMUSG00000022012 | Enox1 | 239188 |
| ENSMUSG00000022054 | Nefm | 18040 |
| ENSMUSG00000022055 | Nefl | 18039 |
| ENSMUSG00000021738 | Atxn7 | 246103 |
| ENSMUSG00000022114 | Spry2 | 24064 |
| ENSMUSG00000035953 | Tmem55b | 219024 |
| ENSMUSG00000035960 | Apex1 | 11792 |
| ENSMUSG00000021871 | Pnp | 18950 |
| ENSMUSG00000025545 | Clybl | 69634 |
| ENSMUSG00000041703 | Zic5 | 65100 |
| ENSMUSG00000036422 | Pcdh8 | 18530 |
| ENSMUSG00000021879 | Dnahc12 | 110083 |
| ENSMUSG00000025545 | Clybl | 69634 |
| ENSMUSG00000047495 | Dlgap2 | 244310 |
| ENSMUSG00000046413 | D230002A01Rik | |
| ENSMUSG00000031734 | Irx3 | 16373 |
| ENSMUSG00000065483 | Mir181c | |
| ENSMUSG00000076338 | Mir181d | |
| ENSMUSG00000058833 | 2810428I15Rik | 66462 |
| ENSMUSG00000007888 | Crlf1 | 12931 |
| ENSMUSG00000062380 | Tubb3 | 22152 |
| ENSMUSG00000074037 | Mc1r | 17199 |
| ENSMUSG00000089704 | Galnt2 | 108148 |
| ENSMUSG00000089704 | Galnt2 | 108148 |
| ENSMUSG00000031930 | Wwp2 | 66894 |
| ENSMUSG00000071064 | Zfp827 | 622675 |
| ENSMUSG00000037940 | Inpp4b | 234515 |
| ENSMUSG00000003573 | Homer3 | 26558 |
| ENSMUSG00000089704 | Galnt2 | 108148 |
| ENSMUSG00000074052 | BC048644 | |
| ENSMUSG00000089704 | Galnt2 | 108148 |
| ENSMUSG00000031557 | Plekha2 | 83436 |
| ENSMUSG00000053399 | Adamts18 | 208936 |
| ENSMUSG00000013033 | Lphn1 | 330814 |
| ENSMUSG00000089704 | Galnt2 | 108148 |

| | | |
|---|---|---|
| ENSMUSG00000031737 | Irx5 | 54352 |
| ENSMUSG00000031834 | Pik3r2 | 18709 |
| ENSMUSG00000080058 | Gm11175 | |
| ENSMUSG00000071076 | Jund | 16478 |
| ENSMUSG00000033009 | Ogfod1 | 270086 |
| ENSMUSG00000085268 | 5033428I22Rik | |
| ENSMUSG00000053560 | Ier2 | 15936 |
| ENSMUSG00000073822 | Gm2529 | |
| ENSMUSG00000055932 | Fto | 26383 |
| ENSMUSG00000033282 | Rpgrip1l | 244585 |
| ENSMUSG00000054823 | Whsc1l1 | 234135 |
| ENSMUSG00000003228 | Grk5 | 14773 |
| ENSMUSG00000006456 | Rbm14 | 56275 |
| ENSMUSG00000087183 | E030007J07Rik | |
| ENSMUSG00000035342 | Lzts2 | 226154 |
| ENSMUSG00000056829 | Foxb2 | 14240 |
| ENSMUSG00000025066 | 6330577E15Rik | 67788 |
| ENSMUSG00000025217 | Btrc | 12234 |
| ENSMUSG00000024986 | Hhex | 15242 |
| ENSMUSG00000024902 | Mrpl11 | 66419 |
| ENSMUSG00000024993 | Fam45a | 67894 |
| ENSMUSG00000004231 | Pax2 | 18504 |
| ENSMUSG00000034336 | Ina | 226180 |
| ENSMUSG00000087828 | U7 | |
| ENSMUSG00000024949 | Sf1 | 22668 |
| ENSMUSG00000024660 | Incenp | 16319 |
| ENSMUSG00000024789 | Jak2 | 16452 |
| ENSMUSG00000078185 | Chml | 12663 |
| ENSMUSG00000073664 | Nbeal1 | 269198 |
| ENSMUSG00000026568 | Brp44 | 70456 |
| ENSMUSG00000084416 | Rpl10a-ps1 | |
| ENSMUSG00000062510 | Nsl1 | 381318 |
| ENSMUSG00000044708 | Kcnj10 | 16513 |
| ENSMUSG00000026623 | Lpgat1 | 226856 |
| ENSMUSG00000026074 | Map4k4 | 26921 |
| ENSMUSG00000039323 | Igfbp2 | 16008 |
| ENSMUSG00000039377 | Hlx | 15284 |
| ENSMUSG00000065905 | U1 | |
| ENSMUSG00000037461 | Ints7 | 77065 |
| ENSMUSG00000062580 | Timm17a | 21854 |
| ENSMUSG00000038855 | Itpkb | 320404 |
| ENSMUSG00000055676 | Gm5069 | |
| ENSMUSG00000054493 | Gm9947 | |
| ENSMUSG00000041040 | Fam117b | 72750 |
| ENSMUSG00000034343 | Ube2f | 67921 |
| ENSMUSG00000060679 | Mrps9 | 69527 |
| ENSMUSG00000012483 | Rpa3 | 68240 |
| ENSMUSG00000089862 | Gm16039 | |
| ENSMUSG00000055818 | A230083G16Rik | |
| ENSMUSG00000030122 | Ptms | 69202 |
| ENSMUSG00000077093 | AC122520.1 | |
| ENSMUSG00000077115 | AC122520.2 | |
| ENSMUSG00000030223 | Ptpro | 19277 |

| | | |
|---|---|---|
| ENSMUSG00000030303 | Far2 | 330450 |
| ENSMUSG00000030068 | Eif4e3 | 66892 |
| ENSMUSG00000055430 | Nap1l5 | 58243 |
| ENSMUSG00000073209 | Klf14 | 619665 |
| ENSMUSG00000045095 | Magi1 | 14924 |
| ENSMUSG00000056445 | 5730446D14Rik | |
| ENSMUSG00000087658 | Gm15051 | |
| ENSMUSG00000014704 | Hoxa2 | 15399 |
| ENSMUSG00000030126 | Tmcc1 | 330401 |
| ENSMUSG00000029811 | Abp1 | 76507 |
| ENSMUSG00000000248 | Clec2g | 70809 |
| ENSMUSG00000065765 | U6 | |
| ENSMUSG00000030127 | Cops7a | 26894 |
| ENSMUSG00000020817 | Rabep1 | 54189 |
| ENSMUSG00000087373 | Gm15892 | |
| ENSMUSG00000018377 | Vezf1 | 22344 |
| ENSMUSG00000070394 | 1810027O10Rik | 69186 |
| ENSMUSG00000081251 | Gm12164 | |
| ENSMUSG00000076031 | AL645948.1 | |
| ENSMUSG00000006575 | Rundc3a | 51799 |
| ENSMUSG00000091753 | Gm17432 | |
| ENSMUSG00000035152 | Ap2b1 | 71770 |
| ENSMUSG00000018733 | Pex12 | 103737 |
| ENSMUSG00000048616 | Nog | 18121 |
| ENSMUSG00000018379 | Srsf1 | 110809 |
| ENSMUSG00000001105 | Ift20 | 55978 |
| ENSMUSG00000020128 | Vps54 | 245944 |
| ENSMUSG00000039989 | Cbx4 | 12418 |
| ENSMUSG00000087675 | Gm11762 | |
| ENSMUSG00000075528 | Aarsd1 | 73635 |
| ENSMUSG00000034120 | Srsf2 | 20382 |
| ENSMUSG00000020818 | Mfsd11 | 69900 |
| ENSMUSG00000081249 | Gm11517 | |
| ENSMUSG00000018666 | Cbx1 | 12412 |
| ENSMUSG00000091095 | Gm17305 | |
| ENSMUSG00000000049 | Apoh | 11818 |
| ENSMUSG00000005917 | Otx1 | 18423 |
| ENSMUSG00000018677 | Slc25a39 | 68066 |
| ENSMUSG00000078627 | March10 | 632687 |
| ENSMUSG00000085578 | Gm11527 | |
| ENSMUSG00000086015 | 4833417C18Rik | |
| ENSMUSG00000020721 | Helz | 78455 |
| ENSMUSG00000018411 | Mapt | 17762 |
| ENSMUSG00000024251 | Thada | 240174 |
| ENSMUSG00000085544 | A930024N18Rik | |
| ENSMUSG00000055660 | Mettl4 | 76781 |
| ENSMUSG00000024182 | Axin1 | 12005 |
| ENSMUSG00000078821 | Gm17414 | |
| ENSMUSG00000015474 | Ppt2 | 54397 |
| ENSMUSG00000024082 | 2410091C18Rik | 73694 |
| ENSMUSG00000053436 | Mapk14 | 26416 |
| ENSMUSG00000071172 | Srsf3 | 20383 |
| ENSMUSG00000050405 | Tmem151b | 210573 |

| | | |
|---|---|---|
| ENSMUSG00000079563 | Pglyrp2 | 57757 |
| ENSMUSG00000060586 | H2-Eb1 | 14969 |
| ENSMUSG00000032624 | Eml4 | 78798 |
| ENSMUSG00000037321 | Tap1 | 21354 |
| ENSMUSG00000067203 | H2-K2 | |
| ENSMUSG00000024073 | Birc6 | 12211 |
| ENSMUSG00000023902 | Zscan10 | 332221 |
| ENSMUSG00000022893 | Adamts1 | 11504 |
| ENSMUSG00000052299 | Ltn1 | 78913 |
| ENSMUSG00000085617 | G730013B05Rik | |
| ENSMUSG00000072214 | Sept05 | 18951 |
| ENSMUSG00000050761 | Gp1bb | 14724 |
| ENSMUSG00000062713 | Sim2 | 20465 |
| ENSMUSG00000022756 | Slc7a4 | 224022 |
| ENSMUSG00000088806 | SNORA84 | |
| ENSMUSG00000022701 | 2610015P09Rik | 212153 |
| ENSMUSG00000022811 | Zfp148 | 22661 |
| ENSMUSG00000090918 | 1700007L15Rik | |
| ENSMUSG00000063445 | Nmral1 | 67824 |
| ENSMUSG00000051483 | Cbr1 | 12408 |
| ENSMUSG00000024317 | Rnf138 | 56515 |
| ENSMUSG00000037416 | Dmxl1 | 240283 |
| ENSMUSG00000024497 | Pou4f3 | 18998 |
| ENSMUSG00000041915 | Ammecr1l | 225339 |
| ENSMUSG00000023036 | Pcdhga11 | 93724 |
| ENSMUSG00000085906 | Gm16882 | |
| ENSMUSG00000023036 | Pcdhga11 | 93724 |
| ENSMUSG00000036299 | BC031181 | 407819 |
| ENSMUSG00000064844 | SNORD58 | |
| ENSMUSG00000062328 | Rpl17 | 100505287 |
| ENSMUSG00000064871 | SNORD58 | |
| ENSMUSG00000064647 | SNORD58 | |
| ENSMUSG00000092097 | Gm5819 | |
| ENSMUSG00000087274 | 1010001N08Rik | |
| ENSMUSG00000005836 | Gata6 | 14465 |
| ENSMUSG00000007440 | Pcdha1 | 12943 |
| ENSMUSG00000056671 | Prelid2 | 77619 |
| ENSMUSG00000024544 | D18Ertd653e | 52662 |
| ENSMUSG00000079614 | Seh1l | 72124 |
| ENSMUSG00000091127 | Gm17238 | |
| ENSMUSG00000074346 | 1700095B22Rik | |
| ENSMUSG00000040896 | Kcnd3 | 56543 |
| ENSMUSG00000064817 | U1 | |
| ENSMUSG00000027868 | Tbx15 | 21384 |
| ENSMUSG00000027985 | Lef1 | 16842 |
| ENSMUSG00000053706 | B430305J03Rik | |
| ENSMUSG00000036894 | Rap2b | 74012 |
| ENSMUSG00000082826 | Gm6097 | |
| ENSMUSG00000038861 | Pi4kb | 107650 |
| ENSMUSG00000044365 | Cxxc4 | 319478 |
| ENSMUSG00000084564 | Mir1895 | |
| ENSMUSG00000027868 | Tbx15 | 21384 |
| ENSMUSG00000028059 | Arhgef2 | 16800 |

| | | |
|---|---|---|
| ENSMUSG00000065773 | U1 | |
| ENSMUSG00000044365 | Cxxc4 | 319478 |
| ENSMUSG00000028195 | Cyr61 | 16007 |
| ENSMUSG00000021288 | Klc1 | 16593 |
| ENSMUSG00000091196 | Gm17330 | |
| ENSMUSG00000036095 | Dgkb | 217480 |
| ENSMUSG00000035653 | Lrfn5 | 238205 |
| ENSMUSG00000020601 | Trib2 | 217410 |
| ENSMUSG00000087421 | Gm5432 | |
| ENSMUSG00000091781 | Gm9590 | |
| ENSMUSG00000054309 | Cpsf3 | 54451 |
| ENSMUSG00000015143 | Actn1 | 109711 |
| ENSMUSG00000005078 | Jkamp | 104771 |
| ENSMUSG00000004151 | Etv1 | 14009 |
| ENSMUSG00000020580 | Rock2 | 19878 |
| ENSMUSG00000021000 | Ctage5 | 217615 |
| ENSMUSG00000035293 | G2e3 | 217558 |
| ENSMUSG00000020952 | Scfd1 | 76983 |
| ENSMUSG00000020607 | Fam84a | 105005 |
| ENSMUSG00000023050 | Map3k12 | 26404 |
| ENSMUSG00000034259 | Exosc4 | 109075 |
| ENSMUSG00000088300 | AC113595.1 | |
| ENSMUSG00000055024 | Ep300 | 328572 |
| ENSMUSG00000080527 | AC160528.1 | |
| ENSMUSG00000022265 | Ank | 11732 |
| ENSMUSG00000075394 | Hoxc4 | 15423 |
| ENSMUSG00000075555 | Gm10863 | |
| ENSMUSG00000037579 | Kcnh3 | 16512 |
| ENSMUSG00000042406 | Atf4 | 11911 |
| ENSMUSG00000037003 | Tenc1 | 209039 |
| ENSMUSG00000062381 | Vps28 | 66914 |
| ENSMUSG00000059323 | Tonsl | 72749 |
| ENSMUSG00000031309 | Rps6ka3 | 110651 |
| ENSMUSG00000081036 | Gm14628 | |
| ENSMUSG00000084053 | Gm14904 | |
| ENSMUSG00000073131 | Vma21 | 67048 |
| ENSMUSG00000083646 | Gm6088 | |
| ENSMUSG00000083493 | Gm15044 | |
| ENSMUSG00000023443 | Esx1 | 13984 |
| ENSMUSG00000079871 | Gm4984 | |
| ENSMUSG00000015291 | Gdi1 | 14567 |
| ENSMUSG00000001962 | Fam50a | 108160 |
| ENSMUSG00000050379 | Sept06 | 56526 |
| ENSMUSG00000071723 | Gspt2 | 14853 |
| ENSMUSG00000050424 | Pnma5 | 385377 |
| ENSMUSG00000079577 | Gm14692 | 666842 |
| ENSMUSG00000042595 | Fam199x | 245622 |
| ENSMUSG00000031167 | Rbm3 | 19652 |
| ENSMUSG00000055188 | 2900002K06Rik | |
| ENSMUSG00000050148 | Ubqln2 | 54609 |
| ENSMUSG00000087271 | 2900008C10Rik | |
| ENSMUSG00000064818 | U6 | |
| ENSMUSG00000050197 | Rhox13 | 73614 |

| | | |
|---|---|---|
| ENSMUSG00000086937 | Gm15063 | |
| ENSMUSG00000077406 | U1 | |
| ENSMUSG00000079508 | Apoo | 68316 |
| ENSMUSG00000037005 | Xpnpep2 | 170745 |
| ENSMUSG00000034480 | Diap2 | 54004 |
| ENSMUSG00000085558 | 4930412C18Rik | |
| ENSMUSG00000049410 | Gm13060 | 100503878 |
| ENSMUSG00000028514 | Usp24 | 329908 |
| ENSMUSG00000042367 | Gjb3 | 14620 |
| ENSMUSG00000086983 | Gm16972 | |
| ENSMUSG00000035969 | Rusc2 | 100213 |
| ENSMUSG00000028524 | Sgip1 | 73094 |
| ENSMUSG00000028830 | AU040320 | 100317 |
| ENSMUSG00000061887 | Ssbp3 | 72475 |
| ENSMUSG00000085980 | Gm12408 | |
| ENSMUSG00000028461 | Ccdc107 | 622404 |
| ENSMUSG00000088088 | RNase_MRP | |
| ENSMUSG00000023286 | Ube2j2 | 140499 |
| ENSMUSG00000028458 | Tesk1 | 21754 |
| ENSMUSG00000048706 | D4Bwg0951e | 52829 |
| ENSMUSG00000041272 | Tox | 252838 |
| ENSMUSG00000084314 | Gm13048 | |
| ENSMUSG00000066026 | Dhrs3 | 20148 |
| ENSMUSG00000087361 | 0610043K17Rik | |
| ENSMUSG00000054679 | Srsf12 | 272009 |
| ENSMUSG00000028849 | Mtap7d1 | 245877 |
| ENSMUSG00000028618 | Tmem59 | 56374 |
| ENSMUSG00000028619 | 2210012G02Rik | 66526 |
| ENSMUSG00000028653 | Trit1 | 66966 |
| ENSMUSG00000028636 | Ppcs | 106564 |
| ENSMUSG00000070806 | Zmynd12 | 332934 |
| ENSMUSG00000066037 | Hnrnpr | 74326 |
| ENSMUSG00000081330 | Gm13013 | |
| ENSMUSG00000048406 | B330016D10Rik | |
| ENSMUSG00000028563 | Tm2d1 | 94043 |
| ENSMUSG00000082144 | Gm12788 | |
| ENSMUSG00000039713 | Plekhg5 | 269608 |
| ENSMUSG00000055296 | D730040F13Rik | 242474 |
| ENSMUSG00000073987 | Ggh | 14590 |
| ENSMUSG00000026988 | Wdsub1 | 72137 |
| ENSMUSG00000010492 | Gm16119 | |
| ENSMUSG00000000823 | Znf512b | 269401 |
| ENSMUSG00000073236 | 2500004C02Rik | |
| ENSMUSG00000042548 | Asxl1 | 228790 |
| ENSMUSG00000001403 | Ube2c | 68612 |
| ENSMUSG00000084571 | n-R5s201 | |
| ENSMUSG00000086131 | BC048594 | |
| ENSMUSG00000039262 | Prrc2b | 227723 |
| ENSMUSG00000008226 | Scrn3 | 74616 |
| ENSMUSG00000080977 | Gm13772 | |
| ENSMUSG00000077999 | AL589870.1 | |
| ENSMUSG00000027490 | E2f1 | 13555 |
| ENSMUSG00000027330 | Cdc25b | 12531 |

| | | |
|---|---|---|
| ENSMUSG00000026883 | Dab2ip | 69601 |
| ENSMUSG00000037143 | 4930529M08Rik | 100503181 |
| ENSMUSG00000075514 | Gm13375 | |
| ENSMUSG00000027011 | Ube2e3 | 22193 |
| ENSMUSG00000027239 | Mdk | 17242 |
| ENSMUSG00000027618 | Nfs1 | 18041 |
| ENSMUSG00000048186 | Bend7 | 209645 |
| ENSMUSG00000087269 | D330023K18Rik | |
| ENSMUSG00000000194 | Gpr107 | 277463 |
| ENSMUSG00000055612 | Cdca7 | 66953 |
| ENSMUSG00000061689 | Dlgap4 | 228836 |
| ENSMUSG00000086638 | 4930405A21Rik | |
| ENSMUSG00000027523 | Gnas | 14683 |
| ENSMUSG00000005505 | Kbtbd4 | 67136 |
| ENSMUSG00000005510 | Ndufs3 | 624814 |
| ENSMUSG00000068267 | Cenpb | 12616 |
| ENSMUSG00000032565 | Nudt16 | 75686 |
| ENSMUSG00000034135 | Sik3 | 70661 |
| ENSMUSG00000031918 | Mtmr2 | 77116 |
| ENSMUSG00000043659 | Npsr1 | 319239 |
| ENSMUSG00000032212 | Sltm | 66660 |
| ENSMUSG00000064921 | U6 | |
| ENSMUSG00000038379 | Ttk | 22137 |
| ENSMUSG00000032368 | Zic1 | 22771 |
| ENSMUSG00000035443 | Thyn1 | 77862 |
| ENSMUSG00000053641 | Dennd4a | 102442 |
| ENSMUSG00000032555 | Topbp1 | 235559 |
| ENSMUSG00000088793 | U6 | |
| ENSMUSG00000060380 | C030014I23Rik | |
| ENSMUSG00000037801 | Iqch | 78250 |
| ENSMUSG00000077428 | SNORA42 | |
| ENSMUSG00000034488 | Edil3 | 13612 |
| ENSMUSG00000051627 | Hist1h1e | 50709 |
| ENSMUSG00000080741 | Gm11398 | |
| ENSMUSG00000078141 | Gm2399 | |
| ENSMUSG00000050295 | Foxc1 | 17300 |
| ENSMUSG00000069309 | Hist1h2an | 665433 |
| ENSMUSG00000069308 | Hist1h2bp | 319188 |
| ENSMUSG00000003992 | Ssbp2 | 66970 |
| ENSMUSG00000086566 | 2810429I04Rik | |
| ENSMUSG00000092091 | 1700066J03Rik | |
| ENSMUSG00000021510 | A530054K11Rik | 212281 |
| ENSMUSG00000021413 | Prpf4b | 19134 |
| ENSMUSG00000042385 | Gzmk | 14945 |
| ENSMUSG00000052957 | Gas1 | 14451 |
| ENSMUSG00000058773 | Hist1h1b | 56702 |
| ENSMUSG00000075032 | Hist1h3i | 360198 |
| ENSMUSG00000001504 | Irx2 | 16372 |
| ENSMUSG00000020027 | Socs2 | 216233 |
| ENSMUSG00000019935 | Slc17a8 | 216227 |
| ENSMUSG00000019878 | Hsf2 | 15500 |
| ENSMUSG00000020069 | Hnrnph3 | 432467 |
| ENSMUSG00000020068 | Pbld2 | 67307 |

| | | |
|---|---|---|
| ENSMUSG00000085666 | Gm9855 | |
| ENSMUSG00000040502 | March09 | 216438 |
| ENSMUSG00000048701 | Ccdc6 | 76551 |
| ENSMUSG00000078429 | Ctdsp2 | 52468 |
| ENSMUSG00000020262 | Adarb1 | 110532 |
| ENSMUSG00000025422 | Agap2 | 216439 |
| ENSMUSG00000048696 | Mex3d | 237400 |
| ENSMUSG00000084658 | AC152062.1 | |
| ENSMUSG00000033255 | Gm5134 | 333669 |
| ENSMUSG00000055657 | E030030I06Rik | 319887 |
| ENSMUSG00000035242 | Oaz1 | 18245 |
| ENSMUSG00000045867 | Cradd | 12905 |
| ENSMUSG00000009681 | Bcr | 110279 |
| ENSMUSG00000058013 | Sept11 | 52398 |
| ENSMUSG00000029381 | Shroom3 | 27428 |
| ENSMUSG00000029535 | Triap1 | 69076 |
| ENSMUSG00000036323 | Srp72 | 66661 |
| ENSMUSG00000063739 | Gm4963 | 243302 |
| ENSMUSG00000029673 | Auts2 | 319974 |
| ENSMUSG00000044092 | C130050O18Rik | 319772 |
| ENSMUSG00000059991 | Nptx2 | 53324 |
| ENSMUSG00000018263 | Tbx5 | 21388 |
| ENSMUSG00000081683 | Fzd10 | 93897 |
| ENSMUSG00000089856 | Fzd10 | |
| ENSMUSG00000029647 | Pan3 | 72587 |
| ENSMUSG00000035456 | Prdm8 | 77630 |
| ENSMUSG00000041697 | Cox6a1 | 12861 |
| ENSMUSG00000045482 | Trrap | 100683 |
| ENSMUSG00000042605 | Atxn2 | 20239 |
| ENSMUSG00000040003 | Magi2 | 50791 |
| ENSMUSG00000014668 | Chfr | 231600 |
| ENSMUSG00000029384 | 2010109A12Rik | 75610 |
