## supplementary table 3 for "A maternal effect regulates global DNA methylation patterns"

**Table 3a.** List of genes containing a hypomethylated region in gene body or promoter region (< 5Kb upstream from TSS) in BALB/c × C57BL/6 compared to C57BL/6 × BALB/c.

| EnsEmbl ID | Gene name | Entrez ID |
|---|---|---|
| ENSMUSG00000047085 | Lrrc4b | 272381 |
| ENSMUSG00000052605 | Isoc2b | 67441 |
| ENSMUSG00000040952 | Rps19 | 20085 |
| ENSMUSG00000043671 | Dpy19l3 | 233115 |
| ENSMUSG00000036915 | Kirrel2 | 243911 |
| ENSMUSG00000078041 | Mir292 | |
| ENSMUSG00000077815 | Mir290 | |
| ENSMUSG00000078008 | Mir291a | |
| ENSMUSG00000078032 | Mir291b | |
| ENSMUSG00000003273 | Car11 | 12348 |
| ENSMUSG00000070802 | Pnmal2 | 434128 |
| ENSMUSG00000048078 | Odz4 | 23966 |
| ENSMUSG00000022103 | Gfra2 | 14586 |
| ENSMUSG00000021975 | Ints9 | 210925 |
| ENSMUSG00000021752 | Kctd6 | 71393 |
| ENSMUSG00000021994 | Wnt5a | 22418 |
| ENSMUSG00000040640 | Erc2 | 238988 |
| ENSMUSG00000022064 | Pibf1 | 52023 |
| ENSMUSG00000091456 | A930011O12Rik | |
| ENSMUSG00000065597 | Mir124a-1 | |
| ENSMUSG00000022114 | Spry2 | 24064 |
| ENSMUSG00000062093 | Gm10110 | 100503296 |
| ENSMUSG00000031842 | Pde4c | 110385 |
| ENSMUSG00000089704 | Galnt2 | 108148 |
| ENSMUSG00000089704 | Galnt2 | 108148 |
| ENSMUSG00000090275 | Cbln1 | |
| ENSMUSG00000031654 | Cbln1 | 12404 |
| ENSMUSG00000091862 | C230036H16Rik | |
| ENSMUSG00000087391 | Gm2694 | |
| ENSMUSG00000063049 | Ing2 | 69260 |
| ENSMUSG00000074357 | AA386476 | 100504975 |
| ENSMUSG00000042812 | Foxf1a | 15227 |
| ENSMUSG00000089704 | Galnt2 | 108148 |
| ENSMUSG00000031883 | Car7 | 12354 |
| ENSMUSG00000047264 | Zfp358 | 140482 |
| ENSMUSG00000031840 | Rab3a | 19339 |
| ENSMUSG00000035559 | Mpv17l2 | 234384 |
| ENSMUSG00000052837 | Junb | 16477 |
| ENSMUSG00000031731 | Ap1g1 | 11765 |
| ENSMUSG00000031530 | Dusp4 | 319520 |
| ENSMUSG00000052085 | Dock8 | 76088 |
| ENSMUSG00000054874 | Pcnxl3 | 104401 |
| ENSMUSG00000004054 | Map3k11 | 26403 |
| ENSMUSG00000024997 | Prdx3 | 11757 |
| ENSMUSG00000044220 | Nkx2-3 | 18089 |
| ENSMUSG00000085432 | A330032B11Rik | |
| ENSMUSG00000038633 | Degs1 | 13244 |
| ENSMUSG00000013997 | Nit1 | 27045 |
| ENSMUSG00000006412 | Pfdn2 | 18637 |

| | | |
|---|---|---|
| ENSMUSG00000026333 | Gin1 | 252876 |
| ENSMUSG00000052423 | B4galt3 | 57370 |
| ENSMUSG00000084799 | Gm11602 | |
| ENSMUSG00000026247 | Ecel1 | 13599 |
| ENSMUSG00000026494 | Kif26b | 269152 |
| ENSMUSG00000026604 | Ptpn14 | 19250 |
| ENSMUSG00000020423 | Btg2 | 12227 |
| ENSMUSG00000051855 | Mest | 17294 |
| ENSMUSG00000029999 | Tgfa | 21802 |
| ENSMUSG00000039419 | Cntnap2 | 66797 |
| ENSMUSG00000064080 | Fbln2 | 14115 |
| ENSMUSG00000034201 | Gas2l1 | 78926 |
| ENSMUSG00000043372 | Hexim2 | 71059 |
| ENSMUSG00000049299 | Trappc1 | 245828 |
| ENSMUSG00000020402 | Vdac1 | 22333 |
| ENSMUSG00000007850 | Hnrnph1 | 59013 |
| ENSMUSG00000003934 | Efnb3 | 13643 |
| ENSMUSG00000051510 | Mafg | 17134 |
| ENSMUSG00000053906 | Mafg | |
| ENSMUSG00000045176 | 2310047M10Rik | 71923 |
| ENSMUSG00000072809 | 9330160F10Rik | |
| ENSMUSG00000002055 | Spag5 | 54141 |
| ENSMUSG00000020346 | Mgat1 | 17308 |
| ENSMUSG00000001510 | Dlx3 | 13393 |
| ENSMUSG00000091968 | Gm17115 | |
| ENSMUSG00000015579 | Nkx2-5 | 18091 |
| ENSMUSG00000015579 | Nkx2-5 | 18091 |
| ENSMUSG00000003198 | Zfp959 | 224893 |
| ENSMUSG00000061607 | Mdc1 | 240087 |
| ENSMUSG00000001525 | Tubb5 | 22154 |
| ENSMUSG00000075229 | Ccdc58 | 381045 |
| ENSMUSG00000022792 | Yars2 | 70120 |
| ENSMUSG00000039830 | Olig2 | 50913 |
| ENSMUSG00000009569 | Mkl2 | 239719 |
| ENSMUSG00000051375 | Pcdh1 | 75599 |
| ENSMUSG00000007440 | Pcdha1 | 12943 |
| ENSMUSG00000024516 | Sec11c | 66286 |
| ENSMUSG00000085906 | Gm16882 | |
| ENSMUSG00000023036 | Pcdhga11 | 93724 |
| ENSMUSG00000023036 | Pcdhga11 | 93724 |
| ENSMUSG00000007440 | Pcdha1 | 12943 |
| ENSMUSG00000024395 | Lims2 | 225341 |
| ENSMUSG00000028093 | Acp6 | 66659 |
| ENSMUSG00000028096 | Gpr89 | 67549 |
| ENSMUSG00000015522 | Arnt | 11863 |
| ENSMUSG00000063052 | Lrrc40 | 67144 |
| ENSMUSG00000028034 | Fubp1 | 51886 |
| ENSMUSG00000043542 | Fam164a | 67306 |
| ENSMUSG00000088484 | SNORA17 | |
| ENSMUSG00000086392 | A330050B17Rik | |
| ENSMUSG00000004127 | Rg9mtd2 | 108943 |
| ENSMUSG00000021171 | Esyt2 | 52635 |
| ENSMUSG00000021294 | Kif26a | 668303 |

| | | |
|---|---|---|
| ENSMUSG00000058669 | Nkx2-9 | 18094 |
| ENSMUSG00000021095 | Gsc | 14836 |
| ENSMUSG00000002900 | Lamb1 | 16777 |
| ENSMUSG00000090980 | Gm17274 | |
| ENSMUSG00000037169 | Mycn | 18109 |
| ENSMUSG00000035653 | Lrfn5 | 238205 |
| ENSMUSG00000091442 | Gm17031 | |
| ENSMUSG00000048118 | Arid4a | 238247 |
| ENSMUSG00000060499 | Rpl10l | 238217 |
| ENSMUSG00000073187 | Gm5784 | |
| ENSMUSG00000061397 | Krt79 | 223917 |
| ENSMUSG00000087446 | 4930578M01Rik | |
| ENSMUSG00000071757 | Zhx2 | 387609 |
| ENSMUSG00000022997 | Wnt1 | 22408 |
| ENSMUSG00000062397 | Zfp706 | 100042023 |
| ENSMUSG00000022332 | Khdrbs3 | 13992 |
| ENSMUSG00000077858 | AL662925.1 | |
| ENSMUSG00000089768 | Tmsb15b1 | 666244 |
| ENSMUSG00000000881 | Dlg3 | 53310 |
| ENSMUSG00000083710 | Gm15078 | |
| ENSMUSG00000068149 | BC049702 | |
| ENSMUSG00000041096 | Tspyl2 | 52808 |
| ENSMUSG00000048007 | Timm8a1 | 30058 |
| ENSMUSG00000001985 | Grik3 | 14807 |
| ENSMUSG00000029061 | Mmp23 | 26561 |
| ENSMUSG00000051351 | Zfp46 | 22704 |
| ENSMUSG00000028664 | Ephb2 | 13844 |
| ENSMUSG00000028655 | Mfsd2a | 76574 |
| ENSMUSG00000086443 | 4933421A08Rik | |
| ENSMUSG00000087413 | Gm11266 | |
| ENSMUSG00000028226 | Mmp16 | 17389 |
| ENSMUSG00000028218 | Fam92a | 68099 |
| ENSMUSG00000083027 | Gm13140 | |
| ENSMUSG00000078766 | Fut9 | |
| ENSMUSG00000028292 | Rars2 | 109093 |
| ENSMUSG00000066148 | Prpf4 | 70052 |
| ENSMUSG00000028830 | AU040320 | 100317 |
| ENSMUSG00000037771 | Slc32a1 | 22348 |
| ENSMUSG00000087563 | Gm14205 | |
| ENSMUSG00000027523 | Gnas | 14683 |
| ENSMUSG00000074743 | Thbd | 21824 |
| ENSMUSG00000027560 | Dok5 | 76829 |
| ENSMUSG00000027523 | Gnas | 14683 |
| ENSMUSG00000026739 | Bmi1 | 12151 |
| ENSMUSG00000051154 | Commd3 | 12238 |
| ENSMUSG00000042631 | Xkr7 | 228787 |
| ENSMUSG00000087600 | Gm14454 | |
| ENSMUSG00000027459 | Fam110a | 73847 |
| ENSMUSG00000027573 | 2310003C23Rik | 76425 |
| ENSMUSG00000027111 | Itga6 | 16403 |
| ENSMUSG00000068859 | Sp9 | 381373 |
| ENSMUSG00000041837 | Pdcd7 | 50996 |
| ENSMUSG00000023495 | Pcbp4 | 59092 |

| | | |
|---|---|---|
| ENSMUSG00000042073 | Abhd14b | 76491 |
| ENSMUSG00000079592 | C1qtnf5 | 235312 |
| ENSMUSG00000034739 | Mfrp | 259172 |
| ENSMUSG00000032350 | Gclc | 14629 |
| ENSMUSG00000034135 | Sik3 | 70661 |
| ENSMUSG00000032368 | Zic1 | 22771 |
| ENSMUSG00000021670 | Hmgcr | 15357 |
| ENSMUSG00000021357 | Exoc2 | 66482 |
| ENSMUSG00000019132 | BC005537 | 79555 |
| ENSMUSG00000052957 | Gas1 | 14451 |
| ENSMUSG00000021596 | Mctp1 | 78771 |
| ENSMUSG00000074798 | Gm10760 | |
| ENSMUSG00000060969 | Irx1 | 16371 |
| ENSMUSG00000038068 | Rnf144b | 218215 |
| ENSMUSG00000033991 | Ttc37 | 218343 |
| ENSMUSG00000006191 | Cdkal1 | 68916 |
| ENSMUSG00000069270 | Hist1h2ac | 665433 |
| ENSMUSG00000088030 | 5S_rR | |
| ENSMUSG00000018102 | Hist1h2bc | 319189 |
| ENSMUSG00000076377 | AC153972.1 | |
| ENSMUSG00000034949 | Zfr2 | 103406 |
| ENSMUSG00000052613 | Pcdh15 | 11994 |
| ENSMUSG00000040345 | Arhgap9 | 216445 |
| ENSMUSG00000019916 | P4ha1 | 18451 |
| ENSMUSG00000005897 | Nr2c1 | 22025 |
| ENSMUSG00000003072 | Atp5d | 66043 |
| ENSMUSG00000019817 | Plagl1 | 22634 |
| ENSMUSG00000003072 | Atp5d | 66043 |
| ENSMUSG00000035754 | Wdr18 | 216156 |
| ENSMUSG00000035745 | Grin3b | 170483 |
| ENSMUSG00000045518 | Onecut3 | 246086 |
| ENSMUSG00000056310 | Tyw1 | 100929 |
| ENSMUSG00000052139 | Bre | 107976 |
| ENSMUSG00000039095 | En2 | 13799 |
| ENSMUSG00000005907 | Pex1 | 71382 |
| ENSMUSG00000029346 | Srrd | 70118 |
| ENSMUSG00000042328 | Hps4 | 192232 |
| ENSMUSG00000090086 | AI480526 | |
| ENSMUSG00000070576 | Mn1 | 433938 |
| ENSMUSG00000039191 | Rbpj | 19664 |
| ENSMUSG00000090531 | Gm17696 | |
| ENSMUSG00000038295 | Atg9b | 213948 |
| ENSMUSG00000029580 | Actb | 11461 |
| ENSMUSG00000053839 | Gm9924 | 791336 |
| ENSMUSG00000053129 | Gsx1 | 14842 |
| ENSMUSG00000042605 | Atxn2 | 20239 |

**Table 3b**. List of genes containing a hypermethylated region in gene body or promoter region (< 5Kb upstream from TSS) in BALB/c × C57BL/6 compared to C57BL/6 × BALB/c.

| EnsEmbl ID | Gene name | Entrez ID |
|---|---|---|
| ENSMUSG00000025477 | Inpp5a | 212111 |
| ENSMUSG00000065368 | U6 | |
| ENSMUSG00000003273 | Car11 | 12348 |
| ENSMUSG00000059824 | Dbp | 13170 |
| ENSMUSG00000049676 | Catsperg1 | 320225 |
| ENSMUSG00000060579 | Fhit | 14198 |
| ENSMUSG00000022209 | Dhrs2 | 71412 |
| ENSMUSG00000054823 | Whsc1l1 | 234135 |
| ENSMUSG00000088176 | 7SK | |
| ENSMUSG00000004637 | Wwox | 80707 |
| ENSMUSG00000089704 | Galnt2 | 108148 |
| ENSMUSG00000071041 | Gm15210 | |
| ENSMUSG00000031562 | Dctd | 320685 |
| ENSMUSG00000052942 | Glis3 | 226075 |
| ENSMUSG00000038949 | Cnst | 226744 |
| ENSMUSG00000026426 | Arl8a | 68724 |
| ENSMUSG00000026504 | Sdccag8 | 76816 |
| ENSMUSG00000029767 | Calu | 12321 |
| ENSMUSG00000029697 | Fezf1 | 73191 |
| ENSMUSG00000035357 | Pdzrn3 | 55983 |
| ENSMUSG00000062519 | Zfp398 | 272347 |
| ENSMUSG00000078651 | Aoc2 | 237940 |
| ENSMUSG00000000282 | Mnt | 17428 |
| ENSMUSG00000039230 | Tbcd | 108903 |
| ENSMUSG00000086475 | Gm12139 | |
| ENSMUSG00000017639 | Rab11fip4 | 268451 |
| ENSMUSG00000087434 | Gm11202 | |
| ENSMUSG00000018698 | Lhx1 | 16869 |
| ENSMUSG00000087211 | 1500016L03Rik | |
| ENSMUSG00000085940 | 4930405D11Rik | |
| ENSMUSG00000018381 | Abi3 | 66610 |
| ENSMUSG00000050860 | Phospho1 | 237928 |
| ENSMUSG00000038805 | Six3 | 20473 |
| ENSMUSG00000023892 | Zfp51 | 22709 |
| ENSMUSG00000073394 | Gm10497 | |
| ENSMUSG00000084025 | Gm6276 | |
| ENSMUSG00000052299 | Ltn1 | 78913 |
| ENSMUSG00000069041 | Slc25a31 | 73333 |
| ENSMUSG00000027883 | Gpsm2 | 76123 |
| ENSMUSG00000081657 | Gm15466 | |
| ENSMUSG00000081594 | Gm15467 | |
| ENSMUSG00000043998 | Mgat2 | 217664 |
| ENSMUSG00000049751 | Rpl36al | 19982 |
| ENSMUSG00000020973 | 1110034A24Rik | 109065 |
| ENSMUSG00000090819 | Gm17603 | |
| ENSMUSG00000054309 | Cpsf3 | 54451 |
| ENSMUSG00000068522 | Aard | 239435 |
| ENSMUSG00000085396 | 6720401G13Rik | |
| ENSMUSG00000082729 | Gm14845 | |
| ENSMUSG00000082756 | Gm14888 | |
| ENSMUSG00000090613 | Gm17630 | |
| ENSMUSG00000073060 | Zxda | |

| | | |
|---|---|---|
| ENSMUSG00000082317 | Gm14933 | |
| ENSMUSG00000084138 | Gm14915 | |
| ENSMUSG00000078020 | AL805935.1 | |
| ENSMUSG00000036572 | Upf3b | 68134 |
| ENSMUSG00000084053 | Gm14904 | |
| ENSMUSG00000028344 | Invs | 16348 |
| ENSMUSG00000062937 | Mtap | 66902 |
| ENSMUSG00000088504 | SNORA17 | |
| ENSMUSG00000040720 | 1110037F02Rik | 66185 |
| ENSMUSG00000086139 | Gm11211 | |
| ENSMUSG00000028223 | Decr1 | 67460 |
| ENSMUSG00000062478 | Ctrc | 76701 |
| ENSMUSG00000037348 | Paqr7 | 71904 |
| ENSMUSG00000028698 | Pik3r3 | 18710 |
| ENSMUSG00000026864 | Hspa5 | 14828 |
| ENSMUSG00000082163 | Gm14276 | |
| ENSMUSG00000027635 | Dsn1 | 66934 |
| ENSMUSG00000081015 | Gm13805 | |
| ENSMUSG00000026805 | Barhl1 | 54422 |
| ENSMUSG00000085087 | Gm13528 | |
| ENSMUSG00000027651 | Rprd1b | 70470 |
| ENSMUSG00000045225 | Olfr1152 | 258103 |
| ENSMUSG00000082056 | Gm13745 | |
| ENSMUSG00000018326 | Ywhab | 54401 |
| ENSMUSG00000032249 | Anp32a | 11737 |
| ENSMUSG00000045594 | Glb1 | 12091 |
| ENSMUSG00000032571 | Pik3r4 | 75669 |
| ENSMUSG00000032220 | Myo1e | 71602 |
| ENSMUSG00000090626 | Tex9 | 21778 |
| ENSMUSG00000032219 | Tex9 | |
| ENSMUSG00000056919 | 4922501C03Rik | 382090 |
| ENSMUSG00000089092 | U6 | |
| ENSMUSG00000021147 | Wdr37 | 207615 |
| ENSMUSG00000091547 | Gm17547 | |
| ENSMUSG00000050334 | C130071C03Rik | |
| ENSMUSG00000061833 | Gm6311 | |
| ENSMUSG00000069171 | Nr2f1 | 13865 |
| ENSMUSG00000038510 | Rpf2 | 67239 |
| ENSMUSG00000083196 | Gm15272 | |
| ENSMUSG00000020023 | Tmcc3 | 319880 |
| ENSMUSG00000045680 | Tcf21 | 21412 |
| ENSMUSG00000029291 | Rufy3 | 52822 |
| ENSMUSG00000029595 | Lhx5 | 16873 |
| ENSMUSG00000085273 | 4930467D21Rik | |
| ENSMUSG00000058503 | Fam133b | 68152 |
| ENSMUSG00000037653 | Kctd8 | 243043 |
| ENSMUSG00000091434 | 1010001B22Rik | |
| ENSMUSG00000066613 | Zfp932 | 69504 |
| ENSMUSG00000029546 | Uncx | 22255 |
| ENSMUSG00000061288 | Taok3 | 330177 |
